# Functional landscape of circular RNAs in human cancer cells

**DOI:** 10.1101/2025.03.24.645016

**Authors:** Peter Hyunwuk Her, Tiantian Li, Ziwei Huang, Xin Xu, Weining Yang, Moliang Chen, Mona Teng, Sujun Chen, Yong Zeng, Stanley Liu, Benjamin Haibe-Kains, Fraser Soares, Jie Ming, Housheng Hansen He

## Abstract

Circular RNAs (circRNAs) constitute a novel class of noncoding RNAs showcasing distinct tissue- and cell-specific expression patterns. Despite the extensive profiling of circRNAs, their individual functions remain poorly understood. To fill this gap, we designed a genome-wide library of 65,300 shRNAs, targeting 9,663 clinically relevant circRNAs and 3,981 of their linear parental genes, and conducted functional screening in seven types of human cancer. We identified a total of 1,342 essential circRNAs (13.9% screened) that impact cell proliferation in at least one cell line, and in 96.5% of the cases, the linear counterparts are not essential. While a shared common subset emerges as functional regulators across all examined cell lines, the majority of circRNAs are functional in a cell type-specific manner. For a comprehensive presentation of the functional circRNA landscape in cancer, we introduce FunCirc, an online database encompassing functional circRNAs across cancer cell lines, coupled with circRNA expression profiles from diverse cancer and tissue types. Our work enhances the understanding of circRNA functions in cancer and provides the scientific community with a resource to further investigate their intricate roles.

## Introduction

Research into cancer biology has traditionally centered around protein-coding genes; however, a recent surge in transcriptomic investigations has turned the spotlight onto other classes of RNA molecules in cancer. Remarkably, more than 90% of the RNAs originating from the human genome fall within the realm of noncoding RNAs (ncRNAs), many of which remain ambiguously defined^1,2^. With recent advances in RNA therapeutics, clinical trials involving ncRNAs are ongoing with the vast majority of ncRNAs being tested as diagnostic or prognostic biomarkers^3^. Classified primarily by size, ncRNAs encompass a diverse array of molecules, including small ncRNAs such as microRNAs, transfer RNAs (tRNAs), PIWI-interacting RNAs (piRNAs), small nucleolar RNAs (snoRNAs) and small nuclear RNAs (snRNAs), and long non-coding RNAs (lncRNAs), which typically exceed 200 nucleotides (nts) in length^4,5^.

Among the myriad of ncRNA classes, circular RNAs (circRNAs) have emerged as a distinct and intriguing group. Generated through a unique back-splicing mechanism, circRNAs exhibit a covalently closed loop structure devoid of polyadenylated tails^4,5^. Although a minority may harbor protein coding potential, the majority function as ncRNAs, modulating gene expression and cellular processes through various mechanisms including interactions with transcription factors, sequestration of microRNAs, and scaffolding proteins^5^. Their circular structure confers enhanced stability and resistance against degradation by exoribonucleases, making them attractive candidates as diagnostic and prognostic biomarkers in cancer^6,7^.

Transcriptomic profiling studies across various cancer cohorts have unveiled dysregulated expression patterns of circRNAs, hinting at their potential significance in tumorigenesis and disease progression^8–11^. In order to assess the functional significance of particular circRNAs, we conducted a pilot loss-of-function analysis using shRNA screens targeting 1,500 most abundantly expressed circRNAs in prostate cancer cell lines and identified 171 circRNAs that influence cell growth^9^. A recent pooled shRNA screening of 3,354 circRNAs identified pathway-linked candidates^12^. RfxCas13d BSJ-gRNA screening for 762 circRNAs demonstrated cell-type-specific functional hits^11^. However, while these pooled RNAi and Cas13-based screens have begun to nominate functional circRNAs, existing efforts have generally been limited in library size, cancer-context breadth, or the ability to disentangle circRNA phenotypes from those of their linear parental transcripts.

Here, we prioritize a list of 9,663 clinically relevant circRNAs for functional RNA interference screening across seven cancer cell types. Our approach identified a core set of circRNAs essential for cell survival across multiple cancer cell lines and highlighted cancer type-specific dependencies. Furthermore, we developed FunCirc, a curated online database integrating genome-scale functional screening and expression of clinically relevant circRNAs. By centralizing functionally validated circRNA data, FunCirc serves as a valuable resource for RNA therapeutics and biomarker discovery, paving the way for more precise cancer treatment strategies.

## Results

### Curation and functional screening of clinically relevant circular RNAs

To identify circRNA for functional screening, we compiled a comprehensive collection of clinically relevant circRNAs sourced from published cohort studies. Notably, oncogenic circRNA profiling data from Vo *et al*.^8^ and Chen *et al*.^9^ contributed mostly to this endeavor. Vo *et al*. uncovered and characterized 232,655 circRNAs in over 2,000 samples across various cancer types through exosome capture RNA sequencing. Chen *et al*. profiled the transcriptomic landscape of circRNAs in localized prostate samples using ultra-deep total RNA-seq on 144 tumors and characterized 76,311 circRNAs. Augmenting these datasets were in-house patient samples from breast (48 samples, 47,759 circRNAs), lung (63 samples, 15,253 circRNAs) and brain cancer (3 samples, 9,899 circRNAs) cohorts, contributing further to the curation of circRNAs for screening (**Figure 1A**).

**Figure 1:**
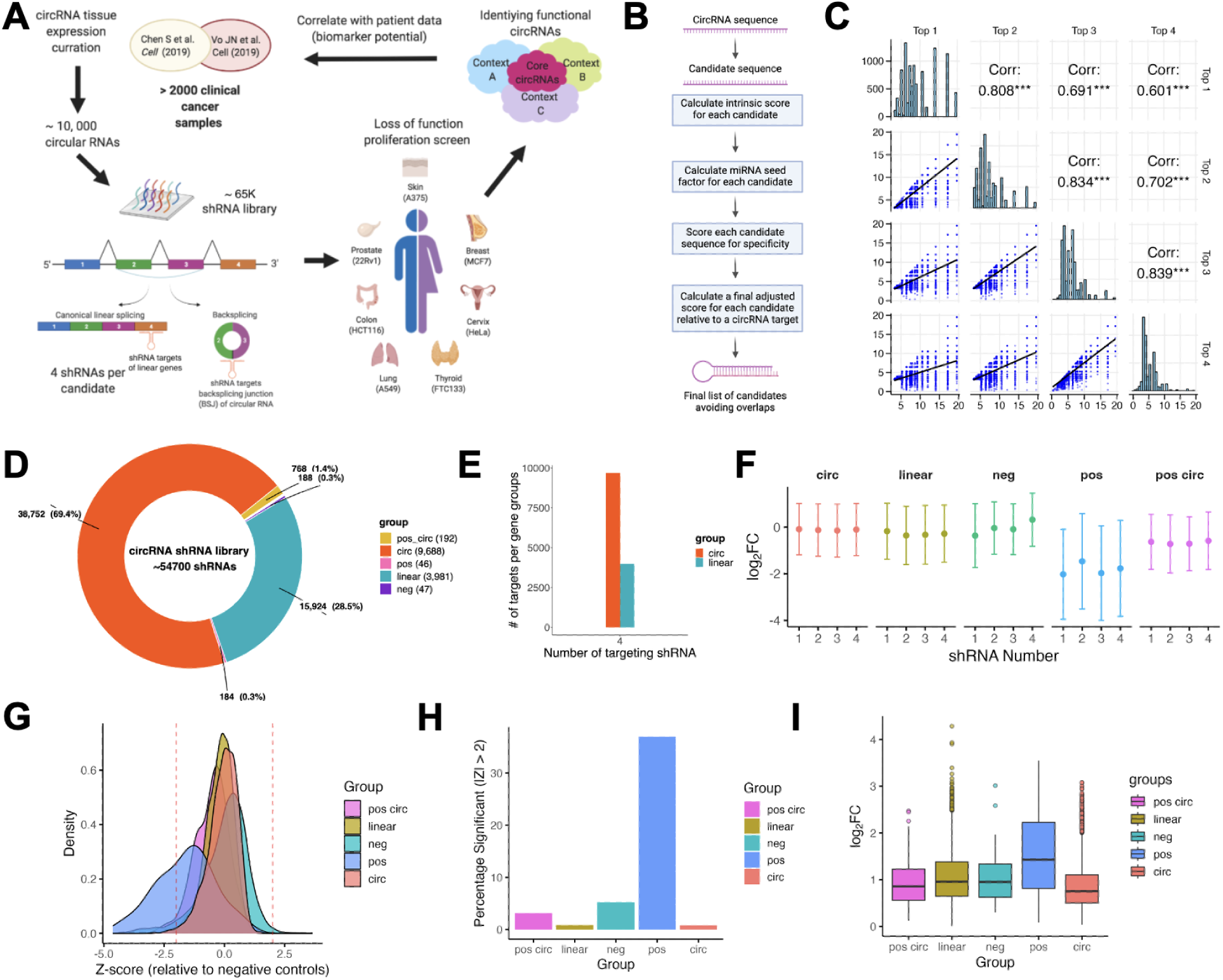
Design and quality control of shRNA screening library targeting cancer associated circRNAs and parental linear transcripts. A) Schematic of pan-cancer shRNA screening. Approximately 10,000 clinically associated circRNAs were filtered from public and internal clinical cohorts, and shRNAs were designed to specifically target the backsplice site of each circRNAs as well as exons of parental linear transcripts. Functional screens were performed in seven major cancer types, including Prostate, Breast, Colon, Lung, Tyroid, Cervix and Skin cancers. B) Schematic of the multi-step shRNA design pipeline for circRNAs C) Pairwise correlation of final adjusted scores among the top four shRNA designs for each circRNA. Scatter plots in the lower off-diagonal cells show the relationship between Top 1–Top 4 ranks, with Pearson’s correlation coefficients (r) and significance (**p < 0.0001) reported in the upper off-diagonal cells. Histograms along the diagonal depict the score distributions for each rank. D) Overview of categories of genes included in the circRNA shRNA screening library. E) Summary of number of shRNAs per linear or circRNA target in the screening library. F) For each group (circ, linear, negative control, positive control, and positive circ control), the mean log_2_ fold change (± SD) is shown for each of the four shRNA positions targeting the same gene or circRNA. Circles represent the average knockdown across all targets in that group for a given position, with vertical bars indicating the standard deviation. G) Each point represents the average log_2_ fold change (LFC) for a given target plotted against its standardized Z-score, where Z = (LFC − mean neg)/SD neg. The dashed horizontal lines mark Z = ±2, commonly used as a significance cutoff. H) Bar chart showing the fraction of shRNAs in each group with |Z| > 2, indicating significant depletion or enrichment relative to negative controls. Positive controls (green) have the highest fraction of “hits” (∼30%). I) Boxplot comparing within-target variability (SD of the log_2_ fold change) for each of the four shRNAs targeting a given gene or circRNA. Dots represent individual genes/circRNAs, boxes denote the interquartile range with the median line, and whiskers extend up to 1.5 × IQR.

Integration of these datasets resulted in 250,380 circRNAs that were filtered based on their expression levels. First, circRNAs were filtered by expression in at least 10% of all samples of each cancer type. Then, they were further filtered by expression in the top 40% of each cancer type. This resulted in an initial list of 18,845 circRNAs for shRNA design, 1,643 of which were overlapping in all cancer types.

We employed a multi-step scoring and filtering process to identify high-confidence shRNA candidates for these circRNAs while minimizing off-target effects (**Figure 1B**, **Methods**). First, we generated candidate sequences by extracting 21-mers across each transcript, skipping the first 25 nt at the 5′ end and the last 150 nt near the 3′ end to avoid poor or highly variable knockdown regions. For circRNAs specifically, we scanned a 34-nt region around the back-splice junction (BJS), capturing 17 nt on each side, and selected 21-mer windows overlapping at least 4 nt of the splice site to ensure circRNA-specific targeting. Each candidate then received an intrinsic score based on a set of rules that penalize or reward features predictive of successful knockdown or undesirable characteristics such as extreme GC content.

Next, we computed a miRNA seed factor to account for potential microRNA-like off-target matches. Candidates with higher predicted seed complementarity to known miRNAs or off-target sites were penalized accordingly, while those showing minimal seed overlap were rewarded. Using the product of the intrinsic score and miRNA seed factor as an intermediate ranking metric, we then assessed sequence specificity via BLASTN searches against the targeted transcriptome. Perfect or near-perfect matches to unintended targets lowered a “specificity factor,” while partial mismatches incurred smaller penalties.

Candidates above an acceptance threshold were further refined by computing a final adjusted score, integrating all intrinsic, seed, and specificity measures. Finally, to avoid overlapping shRNAs that might compete for the same BJS or regulatory regions, we enforced minimal spacing among selected candidates. Only those with acceptable overlap metrics and high adjusted scores advanced to the final library design phase, ensuring each circRNA target was represented by four highly specific shRNAs.

To assess the robustness of our scoring pipeline, we compared the final adjusted scores among the top four shRNA designs (Top 1–Top 4) for each circRNA. The distribution of Top 1 scores was shifted toward higher values compared to Top 2, Top 3, and Top 4, reflecting our rank-based selection criteria (**Figure 1C**). Pairwise correlation analysis among top1–top4 revealed moderate to strong positive correlations (Pearson’s r ranging from 0.60 to 0.80; p < 0.0001), indicating consistent identification of high-scoring shRNAs across multiple ranks. We used a threshold of Adjusted Score > 3 and Adjusted Score > 5 to retain shRNAs with sufficient on-target potency and minimal predicted off-target effects, further refining our candidate list to 9,663 circRNAs. Collectively, these analyses demonstrate that our multi-step pipeline consistently prioritizes the most promising shRNA designs for each circRNA target.

Positive controls such as core essential linear genes and cell line specific essential genes were selected based on previous screening studies^16^ and the DepMap database.

Core essential circRNAs, and cell line specific essential circRNAs were also included from previous studies^8–10^. Negative controls encompassing non-targeting shRNA sequences for GFP, RFP, LacZ, and circRNAs from viral pathogens, as well as non-essential linear genes. The final shRNA library consisted of 65,300 shRNAs, with 4 shRNAs encompassing each target (**Figure 1D**, **E)**.

To assess the functional significance of circRNAs across diverse cancer types on a genome-wide scale, shRNA dropout screens were conducted employing the lentiviral library within 7 cell lines. The cell lines were carefully chosen to represent a broad spectrum of cancer types—including lung (A549), prostate (22Rv1), colorectal (HCT116), melanoma (A375), thyroid (FTC133), breast (MCF7), and cervical (HeLa) cancers, to ensure that our genome-wide shRNA dropout screens capture the functional relevance of circRNAs across diverse tumor contexts. Following puromycin selection, genomic DNA was sampled on Days 0, 8, and 16, and shRNA inserts were PCR amplified for next-generation sequencing (**Figure 1A**). All experiments were performed in triplicates to ensure robustness of the screening.

The shRNA sequencing data were processed using MaGeCK to identify depleted or enriched transcripts. To check for any positional bias in our shRNA designs, we compared the average knockdown across the four shRNA positions for each target. In A549 (**Figure 1F**) and across all cell lines (**Supplemental Figure 1A-F**), there was little difference among positions, indicating minimal positional effects.

We used one-way ANOVA followed by Tukey’s HSD to compare the mean LFC values among the *circ*, *neg*, and *pos* groups across all cell lines. The negative control group exhibited a mean LFC that was 0.019 units higher than that of the circRNA group (diff = +0.01887, 95% CI: [0.00876, 0.02898], padj = 3.61×10⁻⁵), indicating that circRNAs are modestly more depleted than negative controls. In contrast, the positive control group demonstrated a markedly more negative LFC compared to the circRNA group (diff = −1.78130, 95% CI: [−1.88595, −1.67665], padj < 1×10⁻¹⁶) and the negative controls (diff = −1.80017, 95% CI: [−1.90481, −1.69553], padj < 1×10⁻¹⁶). These results suggest that while circRNAs exhibit a moderate knockdown effect relative to negative controls, their depletion is substantially less pronounced than that observed in positive controls, aligning with the concept that circRNAs may modulate cellular processes in a more subtle manner than essential genes.

To assess additional potential off-target knockdowns, we calculated Z-scores for each shRNA relative to the distribution of negative controls. In A549 (**Figure 1G**), and across all cell lines (**Supplemental Figure 1G-L**) the Z-scores for circRNA and linear gene targets form a distribution centered near zero, consistent with the expectation that the majority of circRNAs are non-essential for cell proliferation. Importantly, this distribution is clearly separable from the positive control distribution, which shows marked negative Z-scores reflecting strong depletion of known essential genes. The fact that circRNA-targeting shRNAs do not show inflated variance or systematic shifts relative to negative controls argues against widespread off-target effects. (**Figure 1G**). In contrast the positive-control set displays a clear left-shift (more negative Z-scores), reflecting strong knockdown as expected. Z-scores for circRNA-targeting shRNAs showed markedly lower variability than negative controls in A549 (**Figure 1H**) and across all cell lines (**Supplemental Figure 1M-R**), with only a small percentage exceeding |Z| > 2 compared to negative controls). This tight clustering indicates that the vast majority of circRNA shRNAs produce no detectable effect on cell proliferation, consistent with the expectation that not all circRNAs are non-essential. In contrast, a large number of positive control shRNAs exceeded |Z| > 2, confirming the screen’s sensitivity to detect true essential genes. The lower variability of circRNA shRNAs relative to negative controls further suggests that our multi-step design pipeline successfully minimized off-target effects.

To assess how consistently each target’s four shRNAs produced similar knockdown effects, we calculated the standard deviation (SD) of the log_2_ fold change (LFC) for all targets in each group. In A549 (**Figure 1I**) and across all cell lines **(Supplemental Figure 1S-X)** circRNAs (*circ*) exhibit the lowest median SD, whereas positive controls (*pos*) display the highest variability, with the other groups (*pos circ*, *linear*, and *neg*) falling in between. In A549, Kruskal–Wallis test confirmed significant overall differences among the five groups (Kruskal-Wallis chi-squared = 434.53, *p* < 2.2 × 10⁻¹⁶). Pairwise Dunn tests (Benjamini–Hochberg–adjusted) further indicated that *circ* had significantly lower SD than *linear*, *neg*, and *pos*, while *pos* was significantly more variable than all other groups (*p* < 0.05). This suggests that the four shRNAs targeting each circRNA produce more concordant effects, indicating consistent on-target activity and minimal confounding from variable off-target effects.

To further ensure that our shRNA design pipeline does not inadvertently introduce biases that could affect the screening outcomes, we evaluated the relationship between the final adjusted score and the observed biological effects. Specifically, we computed Pearson’s correlations between the mean final adjusted score (averaged across four shRNAs per circRNA) and both the MAGeCK RRA score and the mean log_2_ fold change. In A549, correlations were statistically significant but negligible in magnitude (r = -0.074, p = 3.5 × 10⁻¹³ for the RRA score; r = -0.077, p = 2.7 × 10⁻¹⁴ for the LFC), explaining less than 1% of variance. The statistical significance reflects the large sample size rather than a meaningful relationship; with nearly 10,000 observations, even trivial effects become statistically detectable. This pattern was consistent across cell lines **(Supplementary Tables 1 and 2)**, confirming that our sequence-based scoring metric does not substantially influence screening outcomes. This lack of meaningful correlation supports the notion that our final adjusted score is not confounded by intrinsic sequence biases and that the functional readouts from the dropout screens reflect genuine biological effects rather than artifacts of shRNA design.

While our multi-step scoring pipeline effectively minimizes off-target effects and maximizes on-target knockdown, several limitations remain. First, in silico specificity checks cannot entirely eliminate the possibility of unanticipated off-targets, particularly in highly repetitive or under-annotated genomic regions. Second, circRNA annotations are still evolving, and our filtering criteria may exclude certain novel or lowly expressed circRNAs. Finally, we focused on 21-mer windows, but alternative shRNA lengths or designs might further optimize knockdown. Therefore, following the screen we sought to validate top circRNA hits with orthogonal methods to confirm both specificity and functional relevance.

### Functional screening identifies known linear dependencies

Following processing using MaGeCK, as shown in 22Rv1 cells, the dropout or enriched levels, as measured by fold change of normalized shRNA abundances, was highly correlated between time points (Day 8 vs. Day 0 and Day 16. vs Day 0) for all the screens (**Figure 2A**, **Supplementary Table 3)**. Furthermore, positive controls showed a significantly higher dropout score (Mann-Whitney U test = 2.56x10^-32^, CLES = 0.21), and negative controls showed a significantly lower dropout score (Mann-Whitney U test = 8.61x10^-48^, CLES = 0.55) (**Figure 2B, Supplementary Figure 2A-F)**, compared to all the linear genes, respectively (“linear” group in **Figure 1B**). The depleted linear transcripts from our screens demonstrated significant positive correlation with publicly available screening data in corresponding cell lines obtained from DepMap^17^ (**Figure 2C**, **Supplementary Table 4)**.

**Figure 2:**
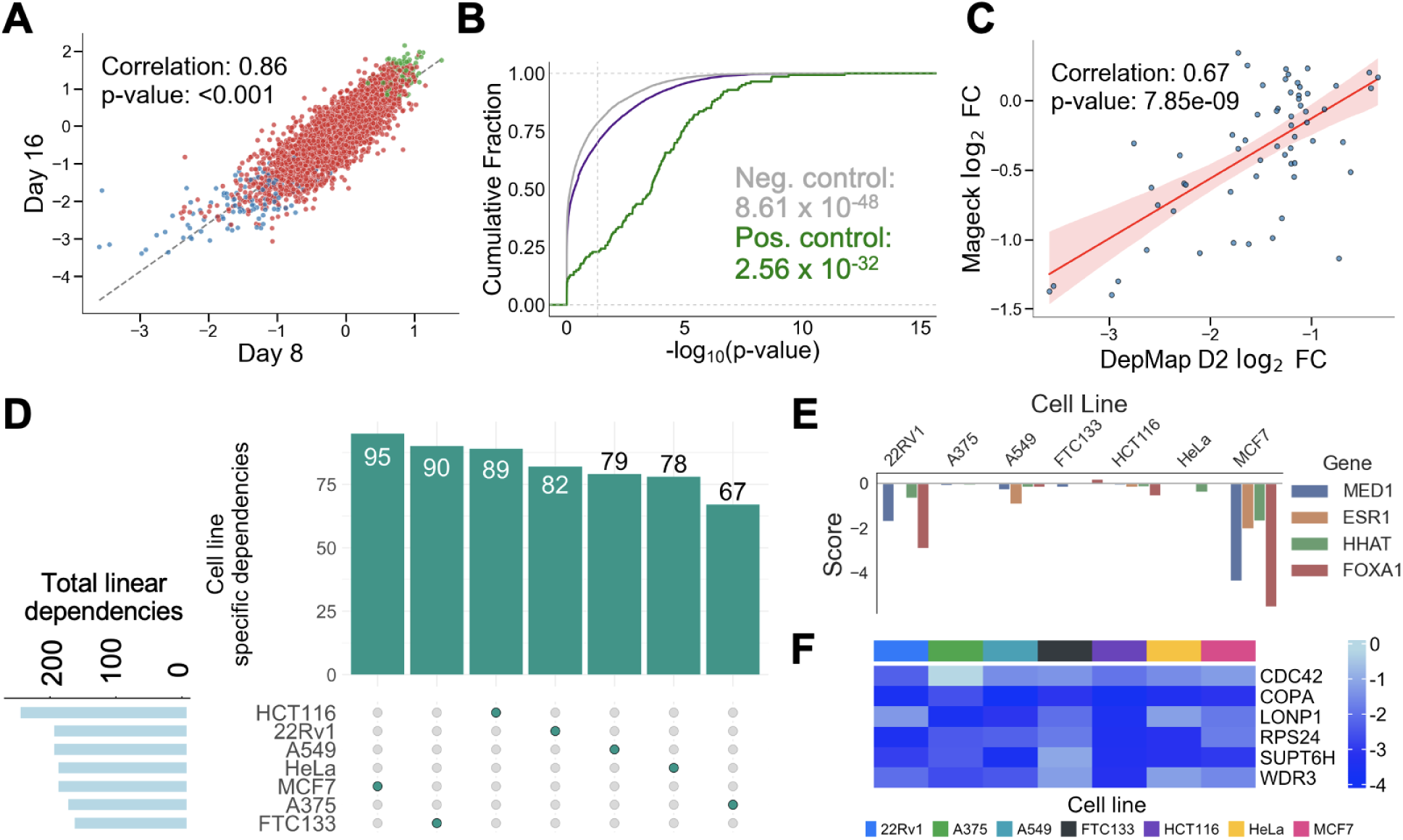
shRNA screens identify common and cancer type specific dependencies of linear transcripts. A) Dropout score measured as log_2_-transformed fold change (log_2_(FC)) in the 22Rv1 cell line of linear transcripts (red), positive controls (green), and negative controls (blue). x and y axes represent comparisons for T8 and T16 samples relative to T0 separately. B) Cumulative distribution of the negative log_10_(p value) for all linear transcripts. Linear transcripts, and positive and negative controls comparing T8 versus T0 samples in the 22Rv1 cell line. p values (Mann–Whitney U test) are relative to the all linear group (purple line). C) Comparison of dropout score in 22Rv1 cells for the union of essential linear transcripts identified from the screens and the DepMap shRNA screening project. D) UpSet plot of cell-line specific and common linear dependencies. Linear transcripts bar graph on the left corner represents the total number of linear dependencies present for each cell line. Columns represent each cell line, with the number of cell-line specific dependencies listed. E) Comparison of dropout score of previously identified linear transcripts with known functional relevance in MCF7. F) Heatmap showing dropout score of common essential linear transcripts identified in our shRNA screens.

We next investigated the cell line-specific and common dropout and enriched linear transcripts identified from our screens. We discovered a total of 491 dropout genes and 363 enriched genes that show significant dependency in at least one cell line. Among the seven cell lines analyzed, HCT116 displayed the highest number of essential dependencies, with 257 identified, while FTC133 showed the lowest, with 173 dependencies (**Figure 2D**). In total, 580 linear genes demonstrated unique dependencies in individual cell lines across cancer types, with the largest subset consisting of 95 dependencies specific to MCF7. Notably, several of these genes, such as MED1, ESR1, HHAT and FOXA1, have been previously reported to be involved in breast cancer biology^18,19,20,21^. Upon analyzing the combined score, which accounts for both the fold change and the statistical significance, MCF7 cell line showed the most substantial changes across these genes, further confirming the prominent role for these genes in breast cancer (**Figure 2E**). A total of 274 genes exhibited dependencies in two or more cell lines, with a set of six genes, including COPA, CDC42, LONP1, SUPT6H, WDR3 and RPS24, depleted across all cell lines (**Figure 2F, Supplementary Figure 3G**). These common essential genes have been previously identified and characterized as oncogenes linked to various cancers ^22,23,24,25,26,27^.

Taken together, our screens identified known cell-type specific and common linear gene dependencies, validating the fidelity of our screens.

### Functional screening identifies core and cell-type specific essential circRNAs

Building on the insights gained from linear transcripts analyses, we utilized MAGeCK to determine the depletion or enrichment of circRNAs in our screens. Similar as observed for linear transcripts, the changes in normalized abundance of shRNAs targeting circRNA back-splicing sites, measured as fold change, showed a strong positive correlation between time points (Day 8 vs Day 0 and Day 16 vs Day 0) in 22Rv1 (correlation = 0.90, p-value = < 0.001) (**Figure 3A**), a pattern consistent across all screens in other cell lines **(Supplementary Table 5)**. Furthermore, positive control circRNAs in 22Rv1 showed a significantly higher dropout compared to all circRNAs ( 4.95x10^-54^, CLES = 0.34), while negative controls showed a significantly lower dropout score (Mann-Whitney U test = 2.28x10^-36^, CLES = 0.54) (**Figure 3B**), a finding replicated in majority of the screens **(Supplementary Figure 2G-L)**.

**Figure 3:**
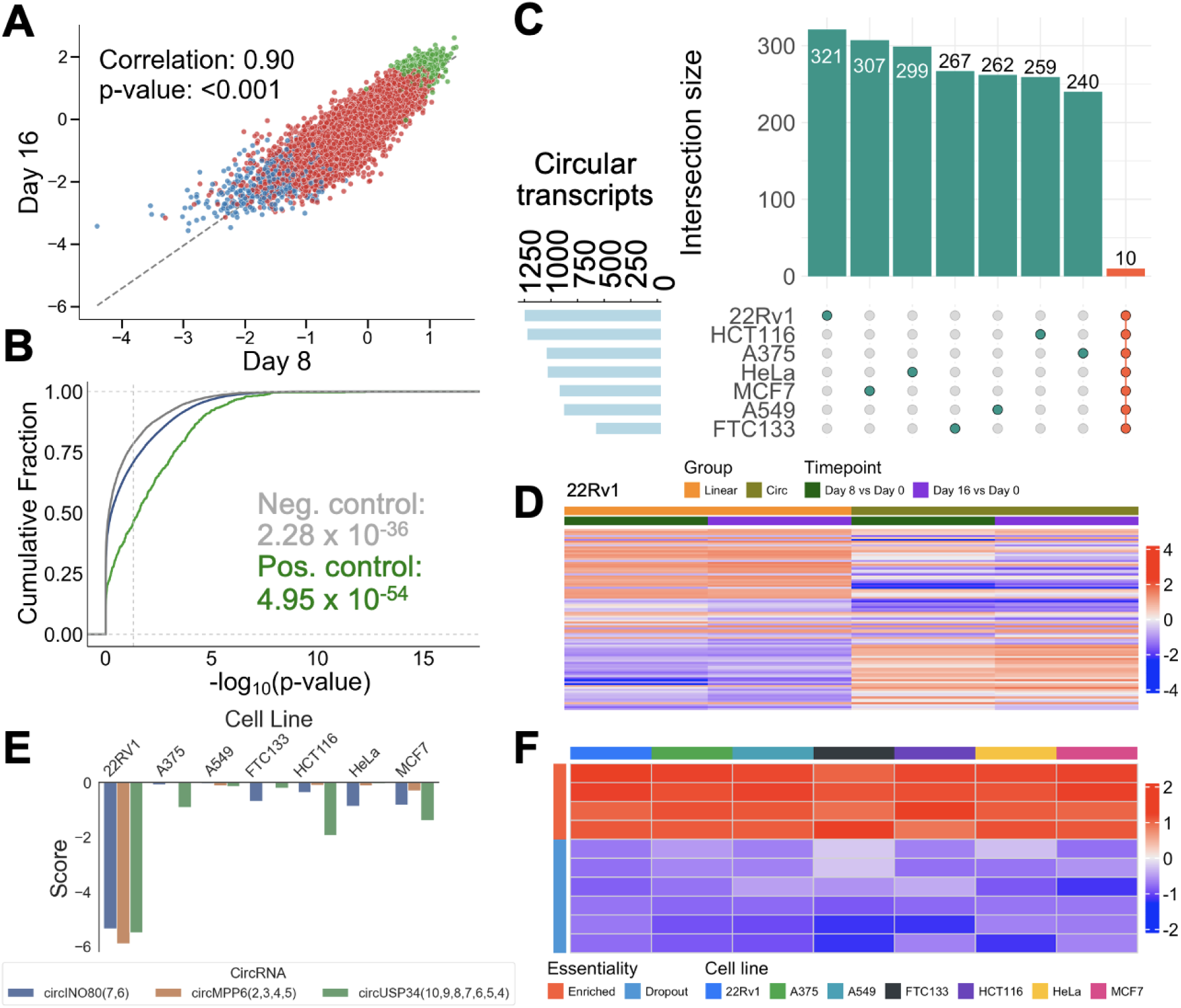
shRNA screens identify common and cancer type specific dependencies of circRNA transcripts. A) Dropout score measured as log_2_(FC) in the 22Rv1 cell line of circular transcripts (red), positive controls (green), and negative controls (blue). x and y axes represent comparisons for T8 and T16 samples relative to T0 separately. B) Cumulative distribution of the negative log_10_(p value) for all circular transcripts. Circular transcripts, and positive and negative controls comparing T8 versus T0 samples in the 22Rv1 cell line. p values (Mann–Whitney U test) are relative to the all circular group (blue line). C) UpSet plot of cell-line specific and common circular dependencies. Circular transcripts bar graph on the left corner represents the total number of circular dependencies present for each cell line. First seven columns (teal) represent each cell line, with the number of functional circular transcripts specific to each cell line listed. The final column (orange) represents the functional circular transcripts shared across all the seven cell lines. D) Depletion of linear and circRNAs following shRNA screening in 22Rv1 cell line. Heatmap shows essential circRNAs along with their linear parental gene. Color represents log_2_(FC) of T8 and T16 compared to T0, with red and blue shades representing negative (depleted) and positive (enriched) values separately. E) Comparison of dropout score of representative functional unknown circRNAs specific to 22Rv1. F) Heatmap showing dropout score of common dropout and enriched circular transcripts identified in our shRNA screens.

Our differential shRNA abundance analysis identified a total of 3,573 circRNAs showing differential dependencies in at least one cell line. The number of differential dependencies ranged broadly, from 571 in FTC133 to 1,198 in 22Rv1 (**Figure 3C**), reflecting the diverse dependence of circRNAs across the cell lines. Importantly, in 96.5% of the cases, circRNA dependencies were distinct from their linear parental genes in at least one cell line (**Figure 3D**, **Supplementary Figure 3A-F).** Among these circRNAs, 1,955 demonstrated unique dependencies in individual cell lines, with prevalence varying from 12-16% across different cell lines (**Figure 3C**). The largest subset, consisting of 331 dependencies, was specific to 22Rv1 cells. Notably, many of these, including cricINO80, circMPP6(2,3,4,5) and circUSP34, with functions unknown in prostate cancer (**Figure 3E**).

While the prevailing trend in circRNA dependencies predominantly displayed functional uniqueness associated with specific cancer cell types, we identified 1,618 circRNAs that showed dependencies in at least two cell lines. Additionally, a core group of circRNAs demonstrated shared functionality across all the seven cancer cell lines screened. This group includes six essential or positive regulators of cell growth (circABHD17B(4,3,2), circGOLIM4(8,7,6), circMTR(3-28), circPSEN1(2,3), circPTK2(7,6,5,4), and circPTK2(7,6,5,4)) and four negative regulators of cell growth (circADGRB3(18,19,20), circARFGEF1(4,3), circCUL1(2-9), and circRacGAP1(17,16,15)) (**Figure 3F**). Importantly, isoforms of circPSEN1, circARFGEF1, circPTK2 and circRACGAP1 align with previously reported functional roles, reinforcing the robustness and reliability of our screening approach.^28–35^.

To evaluate the scope and uniqueness of our circRNA screening library, we compared the circRNAs included in our study with those from previous functional screens. Our library encompassed 9,853 circRNAs and 190 positive control circRNAs, which is significantly more than Li et al.^12^ (762 circRNAs), Liu et al.^36^ (3,354 circRNAs), and Chen et al.^9^ (1,507 circRNAs). After accounting for shared circRNAs, 139 circRNAs were unique to Li et al., 199 to Chen et al., and 1087 to Liu et al., highlighting that our study covers a broader and more comprehensive set of circRNAs (**Supplementary Figure 4A**). This substantial increase in the number of circRNAs screened demonstrates the comprehensiveness of our library, providing a significantly expanded resource for the identification and functional characterization of circRNAs in cancer.

To further assess the utility of our screening, we compared the essential circRNAs identified in our study to those reported by Liu et al., who identified 1,756 circRNAs essential in at least one cell line. Our study identified 1,342 essential circRNAs (plus 78 essential positive control circRNAs) resulting in 1,420 unique circRNAs essential in at least one cell line (**Supplementary Figure 4B**). Of these, 1,220 were novel and did not overlap with those identified by Liu et al. Notably, 200 circRNAs were shared between the two studies. Furthermore, we observed positive correlations in the log_2_ fold changes of overlapping circRNAs screened in our A375 cell line with the skin cell lines from Liu et al. (r = 0.23, p = 2.05 × 10^-3^) (**Supplementary Figure 4C**). This overlap, along with the identification of novel candidates, underscores the utility of our screen in expanding the understanding of circRNA functionality across diverse cellular contexts.

We also compared our results with those from Li et al., who utilized CRISPR-Cas13 screen to identify essential circRNAs in three different cell lines: HT29 (67 circRNAs), 293FT (62 circRNAs), and HeLa (63 circRNAs). The overlaps from this comparison were minimal, with most circRNAs being unique to our screening study (**Supplementary Figure 4D**). Specifically, only 10 essential circRNAs were shared with HeLa, two with 293FT, and one across all studies, highlighting the expanded coverage of our methodology and the unique contributions in identifying additional circRNAs that may have been missed by previous targeted screens.

When compared to our previous work in prostate cancer, 76 of the previous 171 essential circRNAs identified by Chen et al. overlapped with our essential circRNAs (**Supplementary Figure 4E)**. In addition, in the 22Rv1 prostate cancer cell line, our findings correlated with those from Chen et al. (r = 0.25, p = 1.19 × 10^-3^) (**Supplementary Figure 4F**). Collectively, these comparisons with previous studies not only validate our screening approach but also emphasize the novelty and utility of our work in expanding the known landscape of functional circRNAs in cancer.

### Expression does not predict circRNA functional importance in prostate cancer clinical cohorts

We next investigated whether expression could predict circRNA functionality. Because our clinical circRNA profiling was most comprehensive for prostate cancer, we focused this analysis on circRNAs screened in the 22Rv1 prostate cancer cell line. We validated our findings across two independent prostate cancer cohorts, the Chinese Prostate Cancer Genome and Epigenome Atlas^37^ (CPGEA, n = 134 tumors, 134 matched normals) and the Canadian Prostate Cancer Genome Project^38^ (CPCG, n = 145 tumors).

In CPGEA, functional circRNAs (essential and tumor-suppressor combined) showed marginally higher detection rates than non-functional circRNAs (median 26.8% vs 24.6%; Wilcoxon p=0.005). This observation replicated in CPCG (median 27.2% vs 24.8%; Wilcoxon p=0.009) (**Figure 4A**). However, effect sizes were negligible in both cohorts (Cohen’s d=0.075 and 0.060, respectively), and meta-analysis confirmed a pooled effect of only d=0.067 (95% CI: 0.047–0.088) with no heterogeneity (I²=0%, Cochran’s Q p=0.47). Detection patterns were highly correlated across cohorts (Spearman r=0.88; **Figure 4B**), confirming that these observations reflect consistent biology rather than cohort-specific technical artifacts.

**Figure 4:**
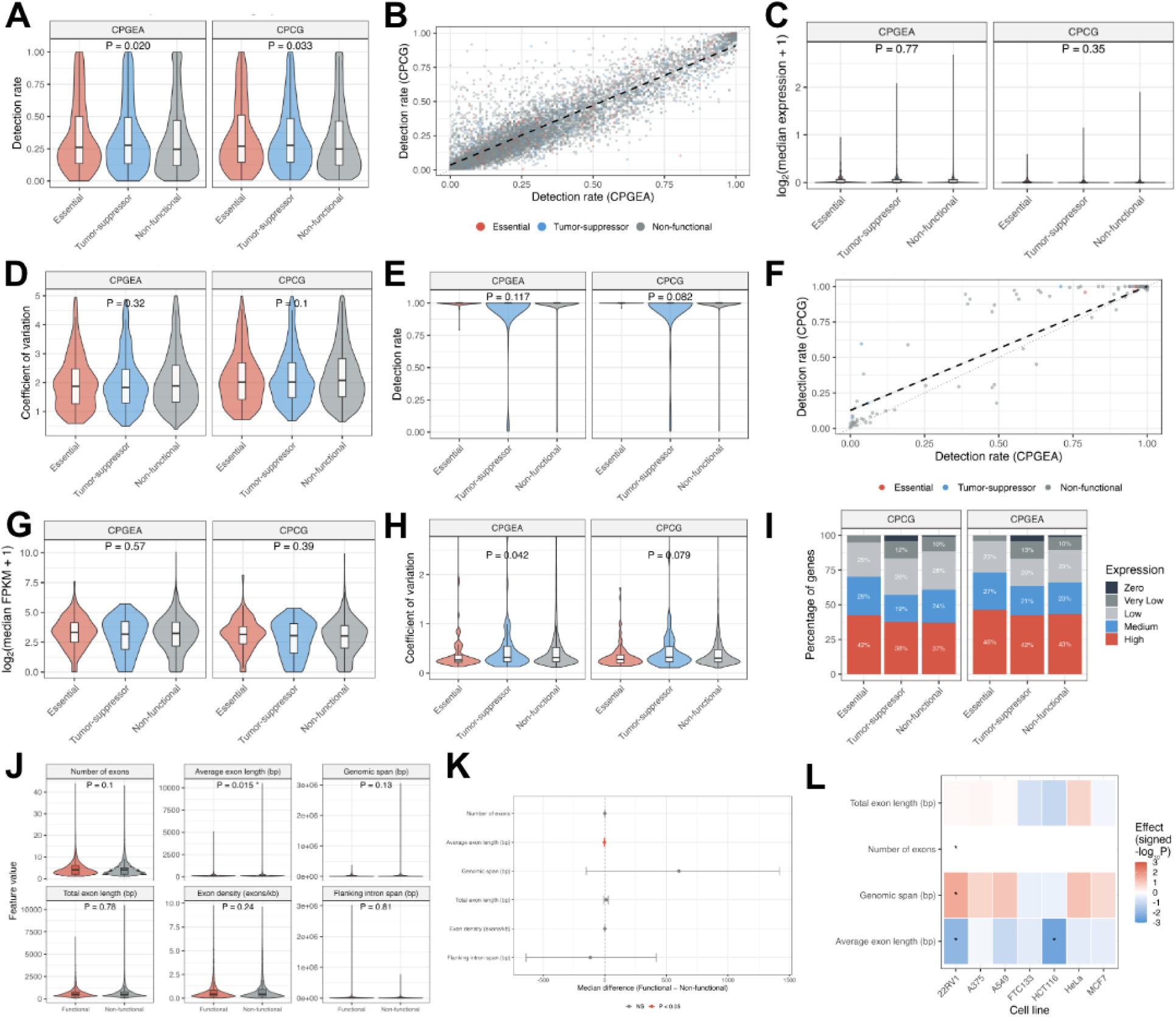
Expression abundance and genomic architecture do not predict circRNA or linear gene functionality. A) Detection rate distributions by functional category in two independent prostate cancer cohorts for circRNAs. Violin plots with embedded box plots show the proportion of patients in which each circRNA was detected (read count > 0) for essential (red), tumor-suppressor (blue), and non-functional (gray) circRNAs in CPGEA (n = 134 patients; left) and CPCG (n = 145 patients; right). B) Cross-cohort reproducibility of detection patterns for circRNAs. Each point represents a circRNA, with detection rate in CPGEA (x-axis) plotted against detection rate in CPCG (y-axis). C) Expression levels among detected circRNAs for circRNAs. Analysis restricted to circRNAs with detection rates >10%. Violin plots show log_2_-transformed median expression values across functional categories. D) Expression stability across patients, measured as coefficient of variation (CV) for circRNAs. Lower CV indicates more consistent expression across the patient population. E) Detection rate distributions by functional category for linear parental transcripts in two independent prostate cancer cohorts. Violin plots with embedded box plots show the proportion of patients in which each gene was detected for essential (red), tumor-suppressor (blue), and non-functional (gray) genes in CPGEA (n = 134 patients; left) and CPCG (n = 146 patients; right). F) Cross-cohort reproducibility of detection patterns for linear parental transcripts. Each point represents a gene, with detection rate in CPGEA (x-axis) plotted against detection rate in CPCG (y-axis). G) Expression levels among detected genes for linear parental transcripts. Analysis restricted to genes with detection rates >10%. Violin plots show log_2_-transformed median FPKM values across functional categories. H) Expression stability across patients, measured as coefficient of variation (CV) for linear parental transcripts. Lower CV indicates more consistent expression across the patient population. I) Expression category distribution across functional classes for linear parental transcripts. Stacked bar plots show the proportion of genes in each expression category (Zero: 0 FPKM; Very Low: 0–1 FPKM; Low: 1–5 FPKM; Medium: 5–10 FPKM; High: >10 FPKM) for each functional category in CPGEA (left) and CPCG (right). J) Comparison of six genomic features between functional circRNAs (exhibiting dependencies in ≥1 cell line) and non-functional circRNAs (no dependencies across all seven cell lines). Violin plots show distributions with embedded box plots indicating median and interquartile range. P-values shown are Benjamini-Hochberg adjusted; asterisk indicates adjusted P < 0.05. K) Bootstrap-estimated effect sizes for genomic feature differences between functional and non-functional circRNAs. Points indicate observed median differences (functional minus non-functional); horizontal lines represent 95% confidence intervals from 2,000 bootstrap iterations. Dashed vertical line indicates zero difference. L) Cell line-specific associations for selected genomic features. Heatmap displays the direction and strength of association between each feature and functionality across seven cell lines, shown as signed −log_10_(P-value). Red indicates functional circRNAs have higher values; blue indicates functional circRNAs have lower values. Asterisks denote nominal P < 0.05.

Among circRNAs with detectable expression (>10% detection rate), absolute expression levels were indistinguishable between functional categories in both cohorts (CPGEA: Kruskal-Wallis p=0.77; CPCG: p=0.35; **Figure 4C**). Expression variability across patients, quantified as the coefficient of variation, showed statistically significant but small differences (CPGEA: p=0.038; CPCG: p=0.026), with median CV values differing by less than 0.08 units across categories in both cohorts (CPGEA: 1.91–1.99; CPCG: 2.09–2.22; **Figure 4D**).

Together, these results demonstrate that circRNA functionality cannot be reliably inferred from expression data alone, underscoring the value of direct functional screening for identifying biologically important circRNAs.

### Linear parental genes exhibit similar expression-independent functionality in prostate cancer clinical cohorts

We performed analogous analyses on the linear transcripts included in our screen. Unlike circRNAs, linear transcripts showed near-universal detection in clinical tumors. Median detection rates exceeded 99% across all functional categories in both CPGEA and CPCG cohorts, with no significant differences between functional and non-functional genes (CPGEA: Kruskal-Wallis p=0.12; CPCG: p=0.08; **Figure 4E**). Detection patterns were highly consistent across cohorts (Spearman r=0.77; **Figure 4F**), confirming these observations reflect reproducible biology. Meta-analysis across both cohorts yielded a pooled effect size for detection rate differences of d=−0.094 (95% CI: −0.126 to −0.061) with no heterogeneity (I²=0%), indicating a small but consistent tendency for functional genes to have marginally lower detection rates opposite to the pattern observed for circRNAs.

Among expressed genes, median FPKM values were indistinguishable between functional categories in both cohorts (CPGEA: Kruskal-Wallis p=0.57; CPCG: p=0.39; **Figure 4G**). Functional genes showed median expression of 8.96 FPKM compared to 8.44 FPKM for non-functional genes in CPGEA (Wilcoxon p=0.56; rank-biserial r=0.027), with similar results in CPCG (7.93 vs 7.15 FPKM; p=0.52; r=0.029). Expression stability, measured as coefficient of variation across patients, showed marginally significant differences in CPGEA (p=0.042) but not CPCG (p=0.079), with median CV values differing by less than 0.05 units across categories (**Figure 4H**).

The expression distribution across functional categories was also uniform. In CPGEA, 46.4% of essential genes, 43.2% of non-functional genes, and 42.3% of tumor-suppressor genes exhibited high expression (>10 FPKM). This was replicated in CPCG (42.3%, 37.1%, and 37.5%, respectively). Zero or near-zero expression was equally rare across all categories (Essential: 0%; Non-functional: 1.0–1.2%; Tumor-suppressor: 4.2%), with no significant association between expression status and functionality (Fisher’s exact p=0.46 in CPGEA; p=0.42 in CPCG; **Figure 4I**).

Collectively, these results demonstrate that for both circular and linear transcripts, functional importance is decoupled from expression abundance, suggesting that cellular context and molecular interactions, rather than transcript levels alone, determine which RNAs become functionally essential in cancer.

### Genomic architectural features show limited association with circRNA functionality

We next examined whether genomic architectural features could distinguish functional from non-functional circRNAs. Using a consensus classification across seven cell lines, we compared six structural features between the circRNAs exhibiting functional dependencies in at least one cell line and the circRNAs showing no functional phenotype in any cell line.

Among the features examined, average exon length showed a modest but statistically significant difference between functional and non-functional circRNAs. Functional circRNAs exhibited shorter average exon lengths compared to non-functional circRNAs (median: 129.2 vs 131.6 bp; Wilcoxon p=0.003, adjusted p=0.015; **Figure 4J**). Bootstrap resampling confirmed this association, with the 95% confidence interval for the median difference excluding zero (Δ=−2.4 bp, 95% CI:−4.5 to −0.5 bp; p=0.017; Figure **4K**). Notably, this direction of effect was consistent across all seven cell lines, with functional circRNAs showing shorter average exon lengths in each individual screen (**Figure 4L**). However, the absolute magnitude of this difference is small.

The remaining genomic features showed no significant associations with functionality after correction for multiple testing. Genomic span showed a trend toward larger values in functional circRNAs (median: 10,034 vs 9,432 bp; Δ=+602 bp, +6.4%), but this did not reach significance (adjusted p=0.13) and the bootstrap confidence interval included zero (95% CI:−151 to +1,418 bp). Importantly, the direction of this effect was inconsistent across cell lines, with functional circRNAs showing larger genomic spans in five cell lines but smaller spans in two (**Figure 4L**). The number of constituent exons was identical between groups (median: 4 exons each; adjusted p=0.10), and total exon sequence length (498 vs 487 bp; adjusted p=0.78), exon density (0.421 vs 0.426 exons/kb; adjusted p=0.24), and flanking intron span (7,791 vs 7,909 bp; adjusted p=0.81) showed no meaningful differences.

These findings suggest that circRNA function is not primarily determined by intrinsic structural properties. The only reproducible association, shorter average exon length in functional circRNAs, is consistent across all cell lines but too small in magnitude to be biologically informative. Combined with our observation that expression abundance also fails to predict functionality, these results indicate that circRNA function emerges from context-dependent interactions with cellular factors rather than from sequence or structural features alone.

### Functional validation of core dropout and enriched circRNAs

To confirm the functionality of the core circRNA dependencies, we first designed divergent primers targeting back-splice junctions. PCR amplification of cDNA from two representative cell lines 22Rv1 and A549, followed by sanger sequencing, verified the back-splicing sites for nine out of the ten core circRNAs. circSCFD1(7,8,9,10), due to unsuccessful amplification, was excluded from subsequent analyses. Subsequent RNase R treatment showed that these circRNAs are highly resistant to exonuclease mediated degradation, contrasting sharply with the linear gene GAPDH, which demonstrated high sensitivity to the treatment (**Figure 5A**, **B)**.

**Figure 5:**
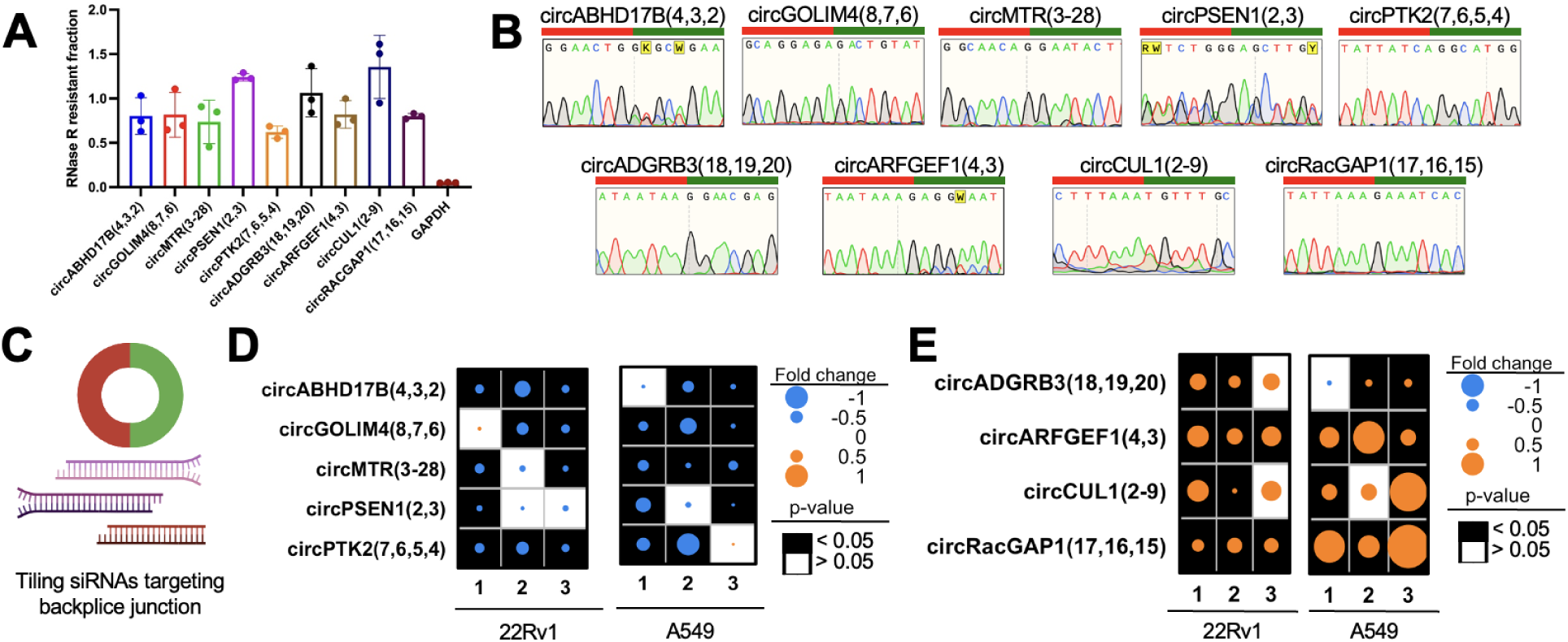
Functional validation of core dropout and enriched circRNAs. A) Core common circRNAs detected by the divergent primers are resistant to RNase R. Data are normalized to their corresponding values in the untreated samples. GAPDH is used as a linear transcript control to demonstrate degradation following RNase R treatment. B) Sanger sequencing of the PCR product by the divergent primers targeting the junction sites of the core common circRNAs to confirm back-splice sequences. The junction between the orange region and the blue region indicates the ‘head-to-tail’ splicing site. C) Schematic of tiling siRNA design targeting the backspliced junction of circRNAs. D-E) Illustration of proliferation assays following circRNA KD in 22Rv1 and A549 cells for the five dropout (D) or four enriched (E) circRNAs. Background shading indicates p value; size and color of dot shows log_2_(FC) of treatment over control. The numbers indicate different siRNA.

To assess the functional role of the core circRNAs, we designed three siRNAs targeting the back splice junction sites for the five core dropout circRNAs and the four enriched circRNAs (**Figure 5C**). We performed cell proliferation assays following the knockdown (KD) of each circRNA in 22Rv1 and A549 cells. Cell proliferation was monitored using the Incucyte live cell imaging system, and fold changes and statistical analyses were done by calculating the area under the curve of confluence measurements over the time course using trapezoidal integration. Upon siRNA-mediated knockdown of these circRNAs, we observed a significant reduction in cell proliferation for the five dropout circRNAs posited to enhance cell survival, consistent with their anticipated biological roles (**Figure 5D**). In contrast, siRNA silencing of the four enriched circRNAs implicated in growth inhibition resulted in a marked increase in proliferation rates in both cell lines (**Figure 5E**). These findings suggest that the core circRNAs play crucial roles in modulating cell growth across different cancer types. This series of experiments not only highlights the functional importance of these core circRNAs, but also validates the reliability of our screening.

### Context specific dependence of circRNA in prostate cancer

In an extension of our validation efforts, we further investigated the function of circRNAs specific to individual cell lines. Notably, circMPP6(2,3,4,5), a previously unreported circRNA, demonstrated dropout exclusively in the 22Rv1 cells. The circular transcript, encompassing 653 nucleotides, is composed of four exons produced by back splicing of exons 2 through 5 (**Figure 6A**). To assess the presence of circMPP6(2,3,4,5) across our panel of screened cell lines, we designed divergent primers targeting this specific back-splice junction. In contrast to the relatively uniform expression of linear MPP6 across cell lines, circMPP6(2,3,4,5) showed highest expression in 22Rv1 cells, underscoring its potential relevance in this cancer type (**Figure 6B**).

**Figure 6:**
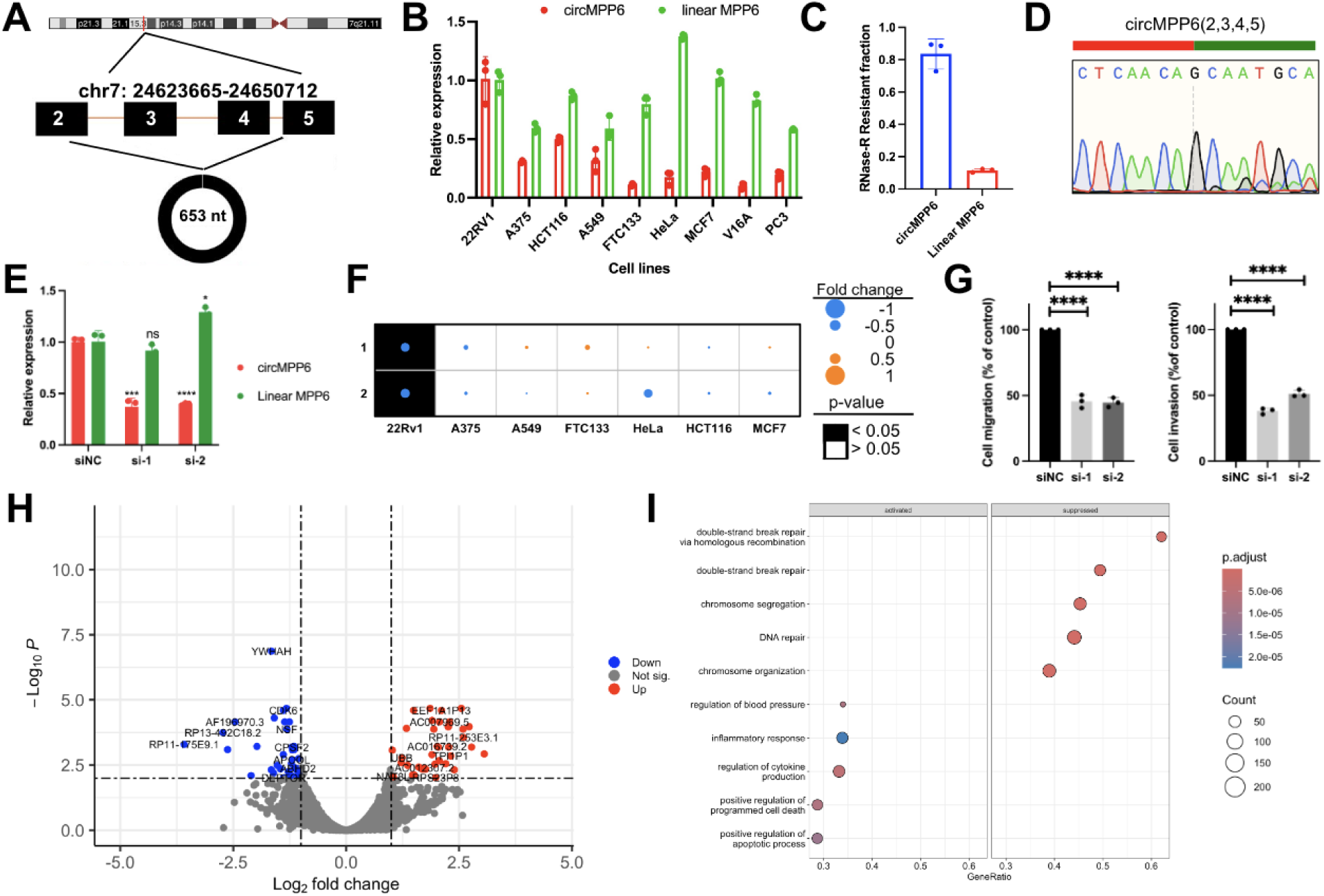
Functional validation of context specific dependence circRNA in prostate cancer. A) Schematic diagram illustrating the formation of circMPP6(2,3,4,5). B) RT-qPCR for circMPP6(2,3,4,5) and its linear parental gene across all cell lines screened. Data are normalized to the 22Rv1 cell line. C) circMPP6(2,3,4,5) detected by the divergent primers is resistant to RNase R. Data are normalized to their corresponding values in the untreated samples. D) Sanger sequencing of the PCR product by the divergent primers targeting the junction sites of circMPP6(2,3,4,5) to confirm back-splice sequences. The junction between the orange region and the blue region indicates the ‘head-to-tail’ splicing site. E) siRNA KD efficiency of circMPP6(2,3,4,5) as determined by RT-qPCR. siNC: Non-targeting siRNA control, si-1, si-2: two tiled siRNAs targeting circMPP6(2,3,4,5). ns, no significance; *, *P* < 0.05; ***, *P* < 0.001; ****, *P* < 0.0001. F) Summary of proliferation assays following circRNA KD across all cell lines for circMPP6(2,3,4,5). G) Migration-invasion assay in 22Rv1 cells following KD of circMPP6(2,3,4,5). NT si: Non-targeting siRNA control, si-1 and si-2: two tiled siRNAs targeting circMPP6(2,3,4,5). ****, *P* < 0.0001. H) Volcano plot showing RNA-seq data in 22Rv1 cells with and without KD of circMPP6(2,3,4,5). The log_2_(FC) indicates the mean differential expression level for each gene contrasting KD and control samples. Grey dots represent non significant (Not sig.) differentially expressed genes between circMPP6(2,3,4,5) KD and control groups, the blue dots represent down-regulated (Down) genes and red dots represent up-regulated genes (Up). I) GO classifications of the differentially expressed genes (log_2_(FC)>1, p<0.05) between circMPP6(2,3,4,5) KD and control groups.

The circular nature of circMPP6(2,3,4,5) was further confirmed through RNAse-R treatment, which significantly reduced the level of linear MPP6, while the circular form demonstrated a high level of resistance to the treatment. Sanger sequencing of the PCR amplicons verified the sequence of the back-splice junction site (**Figure 6C**, **D)**.

To explore the functional significance of circMPP6(2,3,4,5), we designed two siRNAs specifically targeting the back-splicing site. RT-qPCR analysis following siRNA treatment demonstrated an approximate 60% reduction of circMPP6(2,3,4,5) abundance in 22Rv1 cells, with no obvious effect on the level of the linear parental gene (**Figure 6E**), affirming the specificity of these siRNAs. Unlike in other cell lines, where circMPP6(2,3,4,5) knockdown did not yield noticeable impact, the proliferation of 22Rv1 cells was significantly reduced, highlighting a critical, cell line-specific role of circMPP6(2,3,4,5) (**Figure 6F**).

Next, we sought to further characterize the biological functions of circMPP6(2,3,4,5) by performing invasion and migration assays in 22Rv1 cells (**Figure 6G**). Knockdown of circMPP6(2,3,4,5) led to a considerable decrease in these metastatic capabilities, suggesting a role in controlling prostate cancer cell progression.

Further insights into the function of circMPP6(2,3,4,5) were gained through RNA sequencing analysis following circMPP6(2,3,4,5) knockdown. This analysis identified 36 downregulated and 37 upregulated genes (**Figure 6H**). The downregulated genes were significantly enriched in terms associated with genome stability processes such as double-strand break repair via homologous recombination and chromosome organization. In contrast, genes activated by circMPP6(2,3,4,5) knockdown were predominantly linked to regulation of blood pressure, inflammatory response, regulation of cytokine production, and the promotion of programmed cell death and apoptosis (**Figure 6I**).

### FunCirc: a circRNA Function and Clinical Annotation Database

Leveraging data generated from our transcriptome-wide and previously published targeted functional circRNA screens, we developed FunCirc, a comprehensive and open compendium tailored for the scientific community. FunCirc provides clinical annotations and essentiality information for circRNAs across seven major cancer cell types (**Figure 7**). Designed with the needs of the research community in mind, FunCirc offers an extensive collection of over 10,000 circRNAs across multiple studies, enabling detailed exploration of their functions and relationships with their linear counterparts through a user-centric interface (https://funcirc-shiny-153878979048.northamerica-northeast2.run.app/).

**Figure 7:**
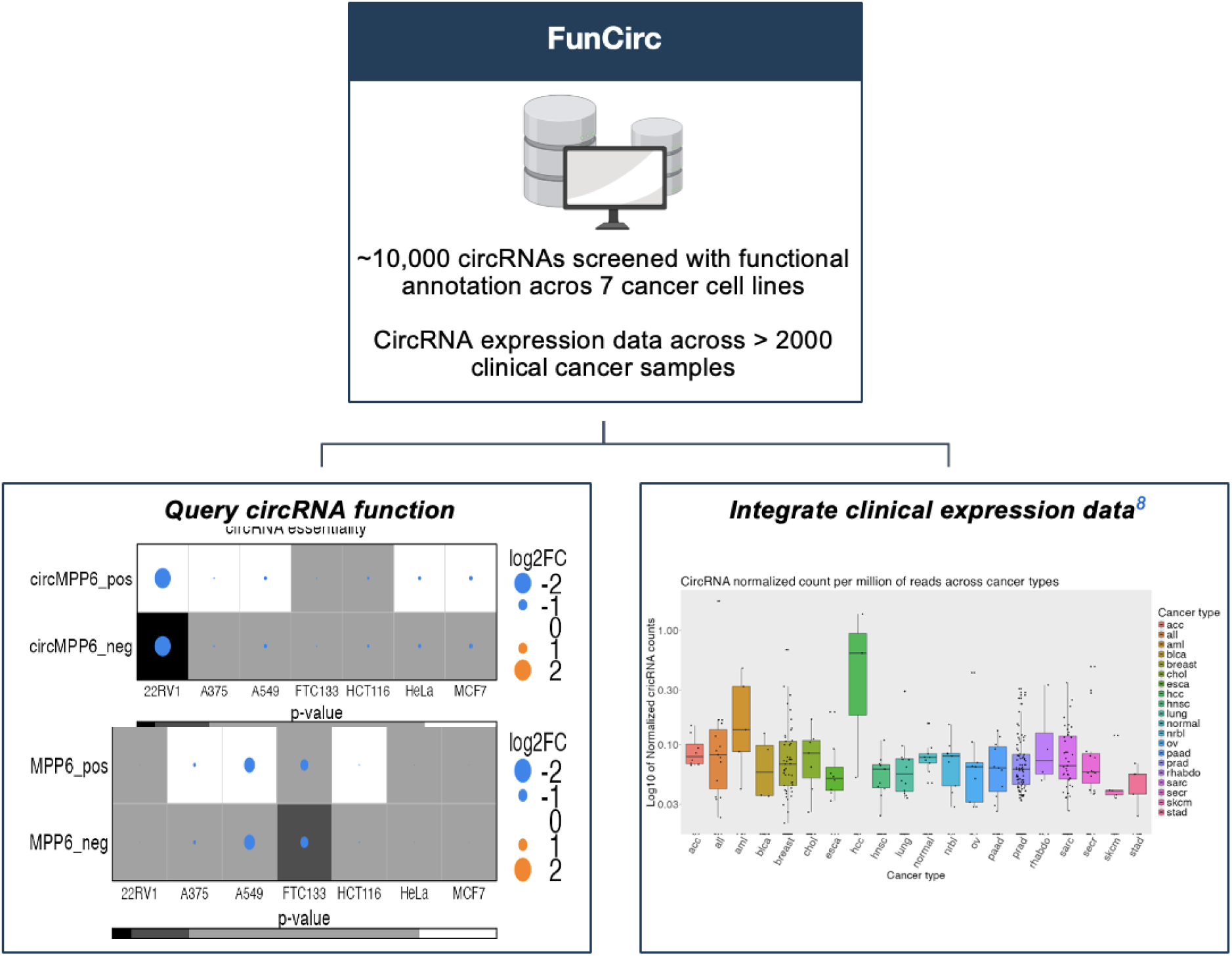
Overview of the FunCirc portal. A schematic overview of the construction of FunCirc (upper panel) and the essentiality and expression outputs (bottom panels).

The platform allows for users to investigate the functional roles of circRNAs across various cell lines and timepoints. CircRNA clinical expression profiles are also integrated and comprise over 2,200 patient tumors and cell lines collected from public studies as well as in-house cohorts^8,9^. This provides a nuanced view of circRNA prevalence across different cancer types, both in cell lines and patient tumours, helping in the identification of potential therapeutic targets and biomarkers.

## Discussion

RNA sequencing technologies have significantly expanded our knowledge of the human transcriptome, revealing the presence of hundreds and thousands of circRNAs. Despite their dynamic expression pattern in various tissues and diseases, including cancer, the functions of these transcripts remain largely unexplored. Majority of circRNAs function as noncoding RNA, and share transcription start sites with their parental linear transcripts, and as a result are not suitable for recently developed CRISPR screening techniques that target protein-coding gene exon DNA or the transcriptional start sites of lncRNAs^39^. Three recent studies^9,11,12^, including one from our group, have utilized pooled shRNA and Cas13 sgRNA designed to specifically target the unique back-splice junctions of circRNAs. However, these efforts have been limited by small library sizes and the number of cell lines used, resulting in an incomplete picture of circRNA functionality.

In this study, we designed over 60,000 shRNAs to target approximately 10,000 cancer-associated circRNAs and their parental linear transcripts, and conducted comprehensive functional screens in cell lines from seven major cancer types. We identified a common group of essential circRNAs across multiple cancer cell lines and hundreds uniquely functional to specific cell lines, some of which have already been documented to play functional roles. Ongoing and future research aimed at elucidating how these shared functions are regulated to execute specific biological actions will deepen our understanding of circRNAs in cancer biology^13,40^.

Our comparisons with previous studies highlight not only the robustness of our screening approach but also the significant contribution to expanding the landscape of functional circRNAs in cancer. The substantial overlap with previous studies validates our findings, while the unique circRNAs identified exclusively in our study suggest new targets for downstream exploration.

Our investigation uncovered hundreds of circRNAs exhibiting specific functions unique to certain cell lines, reinforcing the concept of context-dependent expression of circRNAs^41^. We identified and validated circMPP6(2,3,4,5), as a functional circRNA in prostate cancer. Membrane palmitoylated proteins (MPPs), including MPP6, are involved in crucial biological processes such as cell adhesion and polarity and are often dysregulated in cancers^42,43^. Interestingly, the linear counterpart of MPP6 was associated with prostate cancer progression and was significantly overexpressed in hepatocellular carcinoma^42^. Specifically, the genes exhibiting differential expression were primarily concentrated in processes related to the synthesis of genetic materials, alongside the WNT signaling pathway. Despite some pathway overlaps between the circular and linear target genes, our screening results suggest a unique role for circMPP6(2,3,4,5), distinct from its linear counterpart.

Although shRNA screening offers several advantages, its application to circRNAs presents unique challenges. The design of shRNAs targeting the back splice junctions of circRNAs can lead to overlaps, potentially resulting in skewed or biased screening results. In this study, our robust shRNA pipeline removed several circRNAs because an overlap-free shRNA design was unfeasible. Our guides also showed high concordance for each target. A promising alternative is the use of RNA-targeting type VI CRISPR effectors, known as Cas13 RNases^11,12^. These emergent tools offer a distinct approach for studying circRNAs and could serve as valuable assets for cross-validating experimental outcomes. However, like any evolving technology, the efficacy and reliability of Cas 13 RNases and similar approaches require further development and validation to address their varying extent of efficiency and off-target effects^44,45^.

The increasing interest in circRNA research has led to the development of several circRNA databases such as MiOncoCirc^8^, CircBase^46^, CIRCpedia^47^, circRNAdb^48^, and CSCD^49^. FunCirc stands out as an extensive, cancer-focused resource that integrates functional and clinical data on circRNAs. FunCirc provides fold change values in cellular proliferation across various cancer cell lines post-knockdown of circRNAs and their corresponding linear parental genes. This highlights the unique contributions of circRNAs and also explores the potential interactions between circRNAs and their linear counterparts in cancer progression. Moreover, FunCirc bridges laboratory findings with clinical data, enriching circRNA research.

In summary, we report comprehensive transcriptome-wide circRNA shRNA screens, offering functional annotations across cell lines of diverse cancer types. Our extensive analysis uncovered a group of core circRNAs that positively or negatively regulate cell survival across all cell lines examined as well as hundreds of circRNAs that are crucial for cell survival in a cell-type-specific manner. By validating the functional relevance of circRNAs across various cancer subtypes, our findings underscore their broad impact on cancer biology and the potential impact of circRNAs across varied cancer types. Combining functional and clinical data, our work establishes a robust foundation for future research endeavors, aiming to further elucidate the multifaceted roles of circRNAs in cancer and potentially other diseases.

## Methods

### shRNA library design

Clinically relevant circRNAs for functional validation in cell lines were curated from a list of oncogenic circRNAs compiled from the following published cohort studies: Vo JN et al. 2019, Chen et al. 2019, Wang et al. X, and from additional in-house patient samples [46 Breast cancer cohort (10 normal: 36 tumor) and 3 brain cancer samples). CircRNAs were prioritized based on expression across samples and cancer type. In brief, circRNAs expressed in at least 10% of the samples per cancer type were selected and the top 30-40% expressed in each cancer were selected for library design (9,663 circRNAs). shRNAs were designed for each circRNAs as previously described (Chen et al. 2019). In brief, a 34-nucleotide long sequence around the back splice junction site, with 17 nucleotides on each side, was extracted. A custom script was developed to perform batch shRNA designing with the GPP web portal (https://portals.broadinstitute.org/gpp/public/seq/search). To minimize off-target effects for all candidates, we chose one representative RefSeq transcript and scanned from 25 nt downstream of the coding sequence (CDS) start to 150 nt upstream of the transcript end. We extracted every 21 nt window within this region as a potential short hairpin RNA (shRNA) “candidate.” This exclusion of the extreme 5′ and 3′ regions aims to avoid regulatory elements and regions prone to poor knockdown. Each 21 nt candidate received an intrinsic score derived from rules that either reward or penalize features predictive of efficient knockdown. Factors considered include GC content, presence or absence of specific motifs, and positional attributes (e.g., avoiding extremely close proximity to the CDS boundaries). The sum of these rewards and penalties yields a single numeric score (the “intrinsic factor”) for each candidate. To minimize off-target effects, each candidate was aligned against the entire transcriptome. Candidates exhibiting near-perfect identity (≥ 19 out of 21 nt) to non-target transcripts were heavily penalized or removed altogether. Partial matches (≥ 17 nt) incurred smaller penalties. In certain cases, if a candidate aligned uniquely to a functionally important target (e.g., a known essential gene), a small positive “bonus” factor was applied. The result of this process is a “specificity factor,” which, if it fell below a threshold (e.g., 0.7), caused the candidate to be excluded. The final adjusted score for each candidate was computed by combining (often via multiplication) the intrinsic factor, the specificity factor, and additional penalty or reward terms—such as a miRNA seed factor—to account for potential microRNA-like off-target effects. Candidates with very low specificity or excessively high off-target penalties scored close to zero and were not considered further. In typical linear gene knockdown designs, only the CDS and possibly the 3′ UTR were targeted. Candidates overlapping the CDS–3′ UTR boundary were included if they met length and scoring requirements. For circRNA targets, we additionally scanned across the back-splice junction, ensuring at least 4 nt overlap on each side of the junction to achieve circRNA-specific targeting. Finally, all potential targets were sorted by the adjusted score. For circRNA parental transcripts, the shRNA candidates were scanned from the TRC shRNA library.

The shRNA oligo array was synthesized as 92-mer oligonucleotides (Twist Biosciences), GGAAAGGACGAAACACCGGNNNNNNNNNNNNNNNNNNNNNCTCGAGNNNNNNNNNNNNNNNNNNNNNTTTTTGAATTCTCGACCTCG AGACA (N’s denote shRNA 21nt-target sequence, sense and antisense). Library construction has been previously described (Chen et al., 2019). In brief, the array was amplified by PCR (KAPA HiFi HotStart ReadyMix, KK2601) as a pool using the following primers: TAACTTGAAAGTATTTCGATTTCTTGGCTTTATATATCTTGTGGAAAGGACGAAACAC CGG (Forward) and CCCCCTTTTCTTTTAAAATTGTGGATGAATACTGCCATTTGTCTCGAGGTCGAGAATTC (Reverse). The PCR product was purified and then cloned in the pLKO.1 vector using AgeI/EcoRI (NEB). Ligation was performed using the NEBuilder HiFi DNA Assembly Cloning Kit (NEB, Cat. #E5520S) and transformed into an electrocompetent strain (Endura™ Electrocompetent Cells, 24 rxns (DUOs), Cat. 60242-2). 10-15 Gibson ligation reactions were pooled together to achieve > 300X coverage. Colonies were scraped off plates using LB and plasmid DNA was extracted (NA0310 Sigma GenElute HP Plasmid Maxiprep Kit). The library was submitted for next generation sequencing to confirm adequate library representation of each shRNA.

### shRNA library screening and analysis

Library virus was generated as previously described ^9^ and each cell line was titrated with library virus to achieve a low MOI (∼0.3-0.4). Specifically, MOI was determined by infecting 5-10 million cells with varying amounts of library virus for 24 hours, which were then passed into media with or without 1ug/mL puromycin (ThermoFisher) for 72 hours. A ratio between these two populations was calculated to determine the infection efficiency to achieve a MOI of 0.3. The amount of library virus was scaled up along with the number of cells to ensure that on average every shRNA was represented in 300 cells. For each screen, cells were split into 3 independent batches, passed every 3-4 days, and maintained at 300x coverage for the duration of the screen. A sample was taken from each independent batch on Day 0 (T0), Day 8 (T8) and Day 16 (T16) post-selection for genomic DNA analysis and ∼40-50 million cells were collected per sample. Genomic DNA was extracted (QIAamp Blood Maxi kit, Cat. # 51194) and shRNA inserts were amplified by PCR using standard Illumina next-generation sequencing library preparation kits. The input amount of genomic DNA was calculated to achieve 250x coverage of the library, which was then sequenced on an Illumina HiSeq 2500. Prior to performing screens, all cell lines were authenticated by STR (Geneprint10 panel, Sickkids TCGA) and tested for mycoplasma contamination using the EZ-PCR mycoplasma Test Kit (20-700-20, Biological Industries). All screens were performed with low passage cell lines starting between passages 3-6. Prior to library virus infection, 5 million cells of each cell type were collected for downstream RNA-Seq.

Essential circRNAs were identified from shRNA screen sequencing results with MAGeCK (v0.5.4) software^50^. Sequencing FASTQ files were first trimmed so that the shRNA sequence starts at the first base. The counts of each shRNA in different time points were then counted from the trimmed files with MAGeCK ‘count’ function with default parameters. The Robust Rank Aggregation (RRA) algorithm was applied on the count table. A p value of 0.01 was used as threshold for essentiality.

### Cell culture

MCF7, HeLa, A549, 22Rv1, HCT116, and A375 cell lines were obtained from the American Type Culture Collection (ATCC® CCL-185, CRL-2505, CCL-247, CRL-1619), FTC133 from Sigma (94060901-1VL). 22Rv1 and A549 cells were cultured in RPMI1640 media, HCT116 in McCoy’s 5A media, MCF7, HeLa and FTC133 in DMEM media. All cell culture media was supplemented with 10% FBS (Wisent) and 1% Penicillin and Streptomycin (450-201-EL, Wisent). 293FT cells were cultured in DMEM medium containing 10% FBS (080150, Wisent), L-glutamine (25030-081, ThermoFisher), and non- essential amino acids (11140–050, ThermoFisher) supplemented with 500 μg/mL Geneticin (4727894001, Sigma-Aldrich). All cells were cultured at 37° in 5% CO2.

### Transfection

Control siRNA and all siRNAs targeting linear and circular candidate genes were purchased from MilliporeSigma. Lipofectamine RNAiMax reagent (ThermoFisher; Cat. #13778150) was used to transfect siRNAs. Cells were plated and transfected with the siRNA-lipid complex diluted in Opti-MEM I Reduced Serum Medium (ThermoFisher; Cat. #31985070).

### Cell Proliferation Assays

Cells were seeded in a SARSTEDT 96-well cell culture plate (Cat. # 83.3924) to be 60-80% confluent at transfection. Cells were imaged for 7 days using IncuCyte S3 live cell imaging system (Sartorius, Göttingen, Germany) and the surface area occupied by the cells was calculated, which was then expressed as percent cell confluence. MISSION® siRNA Universal Negative Control #2 (MilliporeSigma) was used as a negative control.

### Invasion Migration Assays

For the migration assay, Corning BioCoat Control Inserts with 8.0µm Pore Polyester (PET) Membrane (Cat # 08-774-162) were used. Cells were pre-treated with siRNAs specific for the circRNA of interest or a non-targeting siRNA for 48 hours. Cells were then trypsinized and resuspended to a concentration of 1*10^5 cells/ml for A549 cells and 2*10^5 cells/ml for 22Rv1. in serum-free RPMI-1640. A total of 500 µl cell suspension was added into the upper chamber, whilst the lower chamber was treated with 750 µl of RPMI-1640 with 10% FBS. Following incubation at 37°C for 20 h, the medium was discarded. Migrated cells were fixed with 100% ethanol for 30 seconds then stained with 0.5% crystal violet at room temperature, washed three times followed by removal of any additional dye with kimwipes and cotton swabs. Representative images were taken under an inverted microscope equipped with a camera.

For the invasion assay, Corning BioCoat Growth Factor Reduced Matrigel Invasion Chamber with 8.0μm Pore Polyester (PET) Membrane (Cat # 08-774-193) were used. Cells were pretreated and trypsinized as discussed above. Prior to adding cell suspension, 500 µl of serum-free RPMI-1640 was added to the transwells and incubated at 37°C for 2 h. The following steps including cell plating, incubation, fixing and staining were conducted as discussed above for the migration assay. Representative images were taken under an inverted microscope equipped with a camera.

### Real-time PCR

Total RNA was purified with the RNeasy Mini Kit (QIAGEN, Cat. # 74106) and DNA was removed by performing on-column DNAse treatment (QIAGEN, Cat. # 79254). cDNA was reverse transcribed using the High Capacity cDNA Reverse Kit (4368814, Applied Biosystems).

For RNase R treatment, one μg total RNA was incubated for 15 min at 37 °C with or without 10 U RNase R (Epicenter, Cat. #RNR07250,) followed by phenol-chloroform (ThermoFisher, Cat. #15593-049) purification. cDNA was reverse transcribed using the High-Capacity cDNA Reverse Transcription Kit (Applied Biosystems, Cat. #4368814).

Divergent primers to detect circRNAs were designed for PCR products of sizes ranging from 100-150 bp and spanning the junction site. Convergent primers that targeted the parental linear mRNA counterparts were exclusive of the circRNA generating region for the same size range. Expression levels of circRNAs and their parental linear RNAs were quantified using the primers listed in (Table S4) and PowerUp SYBR Green Master Mix (Applied Biosystems, Cat. #A25742,) in StepOnePlus Real-Time PCR System (Applied Biosystems) or CFX96 Touch Real-Time PCR Detection System (Bio-Rad). RPS28 was used as the endogenous control gene. The results of RT-qPCR were analyzed by the 2-ΔΔCT method.

### RNA-seq Mapping and Data processing

RNA was extracted with TRIzol (ThermoFisher, 15596026) and processed with a RNA-seq library preparation kit (Illumina, RS-122-2101) to produce libraries for deep sequencing on Illumina Novaseq X Plus. Library preparation and sequencing were performed according to the manufacturer’s protocol. RNA-seq raw reads were first filtered by trim_galore (v 0.5.0) then mapped to the human genome (hg38) by using STAR (V 2.4.2a) ^51^ software with default parameters. The hg38 GENCODE gene list was used for all transcription level analysis. RNA-seq reads strands were determined by RSeQC (V 2.6.1)^52,53^. HTSeq (0.11.0)^54^ was used to get gene level read counts from STAR mapped bam files. The resultant gene read count table was subjected to DESeq2 (V 1.22.2) ^55^ for differential gene analysis and a cutoff of 0.01 for FDR was chosen to identify significant differential genes. log_2_FoldChange < (-1) and log_2_FoldChange > 1 were chosen as up regulated genes and down regulated genes respectively.

### Data Analysis & Visualization

Data analysis was conducted using R (v4.1.0) and Python (v3.8.18). Data visualization was conducted using ggplot2 (v3.36) ^56^, MAGeCKFlute (v1.6.5) ^50,57^, forestplot (v1.10), VennDiagram (v1.7.3) ^58^, ComplexUpset (v1.3.3)^59^, ComplexHeatmap (v.2.12.0)^60^, BoutrosLab.plotting.general (v. 7.1.0)^61^, Guitar (v2.16.0)^62^, ClusterProfiler (v4.8.3)^63^, DESeq2 (v1.40.2)^55^, AnnotationDbi (v.1.62.2)^64^, DOSE (v.3.26.2)^65^, org.Hs.eg.db (v.3.17.0)^66^, EnhancedVolcano (v1.18.0)^67^, Seaborn (v.0.12.2)^68^, Scipy (v1.10.1)^69^, Matplotlib (v3.7.3)^70^. Figure 1A was created using BioRender.com. Database was created using RShiny^71^.

## Data Availability

The DepMap DEMETER2 Data v6 data was downloaded from the DepMap portal (https://depmap.org/portal/download/all/)^17^. CircRNA data from cancer clinical samples were obtained from the MiOncoCirc database (https://mioncocirc.github.io/)^8^. The Raw sequencing data of shRNA screening and RNA-seq generated in this study can be accessed at NCBI under the accessions GSE269567 (reviewer token: gpadomagbhcdfst).

## Code Availability

All R packages used are available online as described in the method section. Code used to generate the FunCirc database can be found here: https://github.com/HansenHeLab/FunCirc

## Author Contributions

**Designed studies:** P.H., T.L., F.S., M.J. and H.H.H.

**Performed experiments:** P.H., F.S., Z.H., X.X., W.Y., M.T.,

**Data Analysis:** P.H., T.L., M.C., S.C., Y.Z.

**Manuscript First Draft:** P.H., F.S., Z.H., T.L., M.J., and H.H.H.

**Revised & approved manuscript:** All authors

## Acknowledgement

This work was supported by the Princess Margaret Cancer Foundation (886012001223 to H.H.H.), Canadian Cancer Society (TAG2018-2061), CIHR operating grants (142246, 152863, 152864 and 159567 to H.H.H.), Terry Fox New Frontiers Program Project Grant (PPG19-1090 and PPG23-1124 to H.H.H.). H.H.H. holds Tier 1 Canada Research Chair in RNA Medicine. X.X. was supported by the Prostate Cancer Foundation Young Investigator Award (21YOUN06). M.T. was supported by a CIHR Doctoral Award. P.H.H was supported by a MOHCCN Health Informatics Award and a CIHR Doctoral Award.

J.M. was supported by the Science Fund of the National Natural Science Foundation of China (No. 82270830) and Hubei Provincial Science & Technology Innovation Team Grant (No. 2022CFB072).

## Inclusion & Ethics Statement

We actively took measures to prevent stigmatization, incrimination, or discrimination. Safety measures were implemented to protect participants and researchers, with risk management plans established for potential health, safety, and security risks. We also ensured that local and regional research relevant to this study was appropriately cited, with efforts made to promote diverse perspectives in our reference list.

## Supplementary Material

**Supplementary Figure 1:**
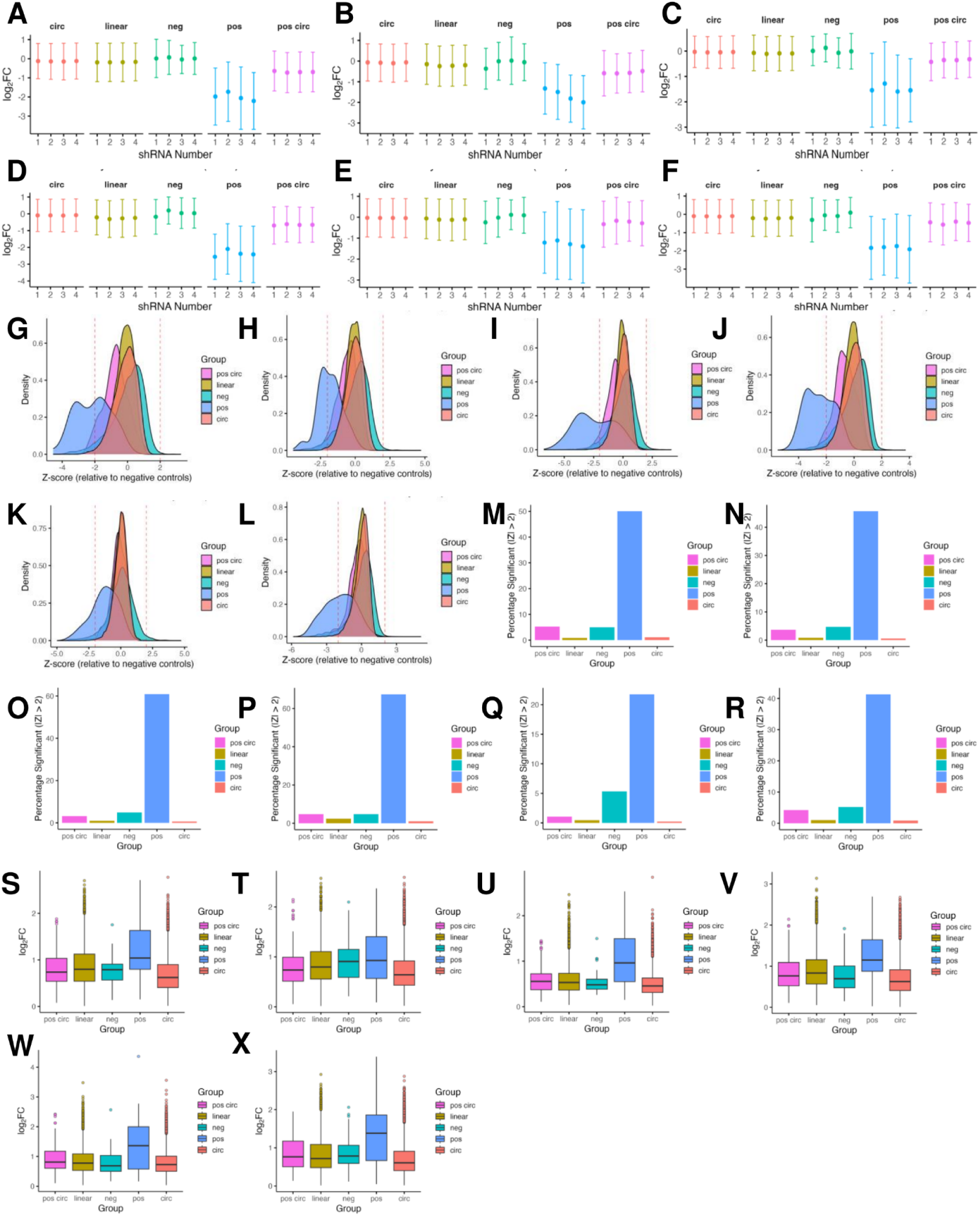
Design and quality control of shRNA screening library targeting cancer associated circRNAs and parental linear transcripts across all cell lines. A-F) (A) 22Rv1, (B) A375, (C) MCF7, (D) HCT116, (E) FTC133, (F) HeLa. For each group (circ, linear, negative control, positive control, and positive circ control), the mean log_2_ fold change (± SD) is shown for each of the four shRNA positions targeting the same gene or circRNA. Circles represent the average knockdown across all targets in that group for a given position, with vertical bars indicating the standard deviation. G-L) (G) 22Rv1, (H) A375, (I) MCF7, (J) HCT116, (K) FTC133, (L) HeLa. Each point represents the average log_2_ fold change (LFC) for a given target plotted against its standardized Z-score, where Z = (LFC − mean neg)/SD neg. The dashed horizontal lines mark Z = ±2, commonly used as a significance cutoff. M-R) (M) 22Rv1, (N) A375, (O) MCF7, (P) HCT116, (Q) FTC133, (R) HeLa. Bar chart showing the fraction of shRNAs in each group with |Z| > 2, indicating significant depletion or enrichment relative to negative controls. Positive controls (green) have the highest fraction of “hits” (∼30%). S-X) (S) 22Rv1 (Kruskal-Wallis chi-squared = 563.63, p-value < 0.001), (T) A375 (Kruskal-Wallis chi-squared = 440.48, p-value <0.001), (U) MCF7 (Kruskal-Wallis chi-squared = 277.96, p-value <0.001), (V) HCT116 (Kruskal-Wallis chi-squared = 680.35, p-value <0.001), (W) FTC133 (Kruskal-Wallis chi-squared = 73.24, p-value <0.001), (X) HeLa (Kruskal-Wallis chi-squared = 258.94, p-value <0.001). Boxplot comparing within-target variability (SD of the log_2_ fold change) for each of the four shRNAs targeting a given gene or circRNA. Dots represent individual genes/circRNAs, boxes denote the interquartile range with the median line, and whiskers extend up to 1.5 × IQR.

**Supplementary Figure 2:**
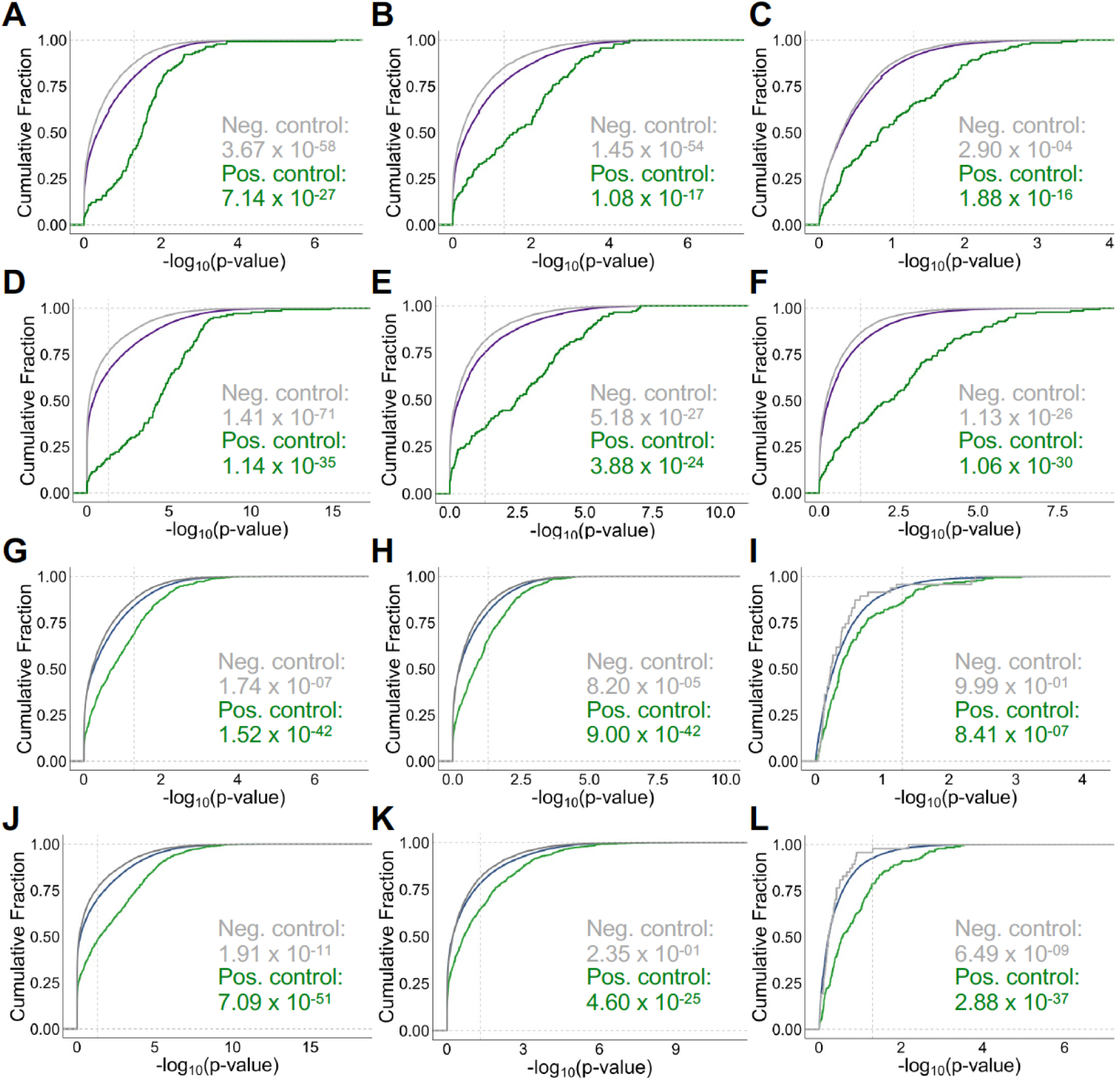
Quality measurement of shRNA screens of circular and linear transcripts. A-F) Cumulative distribution of the negative log_10_ transformed p-value for all linear transcripts, positive and negative controls comparing Day 8 versus Day 0 samples in A375, A549, FTC133, HCT116, HeLa, MCF7 cell lines. One-sided Mann-Whitney U test was used to compare groups to the linear group (purple line). CLES between the linear and positive control group are 0.24, 0.29, 0.30, 0.20, 0.25 and 0.22 respectively. CLES between the linear and negative control group are 0.56, 0.56, 0.51, 0.56, 0.54 and 0.54 respectively. G-L) Cumulative distribution of the negative log10 transformed P value for all circular transcripts, positive and negative controls comparing Day 8 versus Day 0 samples in A375, A549, FTC133, HCT116, HeLa, MCF7 cell lines. One-sided Mann-Whitney U test was used to compare groups to the linear group (purple line). CLES between the linear and positive control group are 0.36, 0.36, 0.44, 0.34, 0.39 and 0.37 respectively. CLES between the linear and negative control group are 0.52, 0.51, 0.49, 0.52, 0.50 and 0.52 respectively.

**Supplementary Figure 3:**
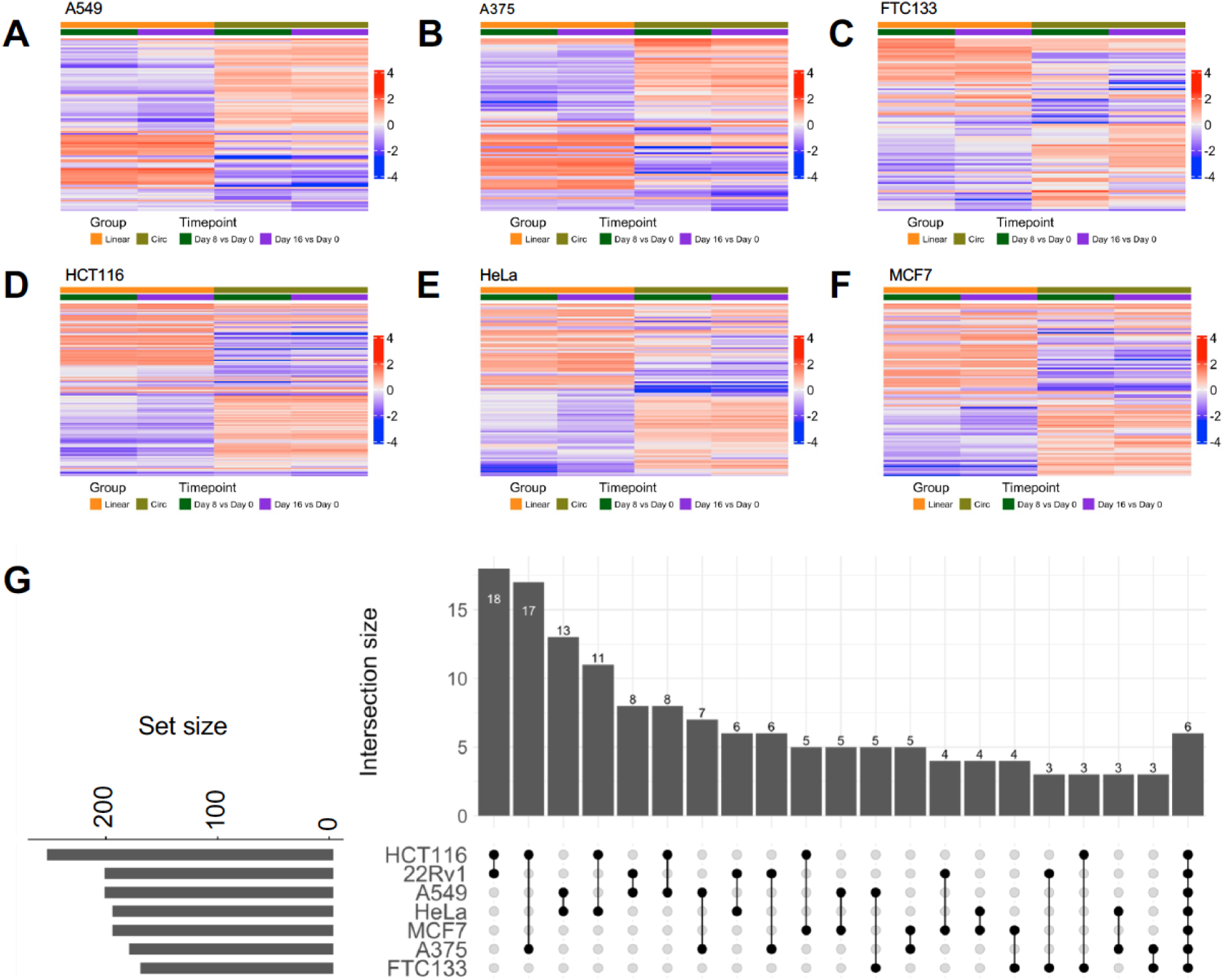
circular and linear dependencies identified in the shRNA screens. A-F) Depletion of linear and circular RNAs following shRNA screening in A549 (A), A375 (B), FTC133 (C), HCT116 (D), HeLa (E) and MCF7 (F) cell lines. Heatmap shows essential circRNAs along with their linear parental gene. Color represents log_2_(FC) of T8 and T16 compared to T0, with red and blue shades representing negative (depleted) and positive hits. G) UpSet plot of linear dependencies common for two sets of cell lines along with the set common across cell lines. Linear transcripts bar graph on the left corner represents the total number of linear dependencies present for each cell line. Columns represent each set of cell lines, with the number of transcripts listed in or above the column.

**Supplementary Figure 4:**
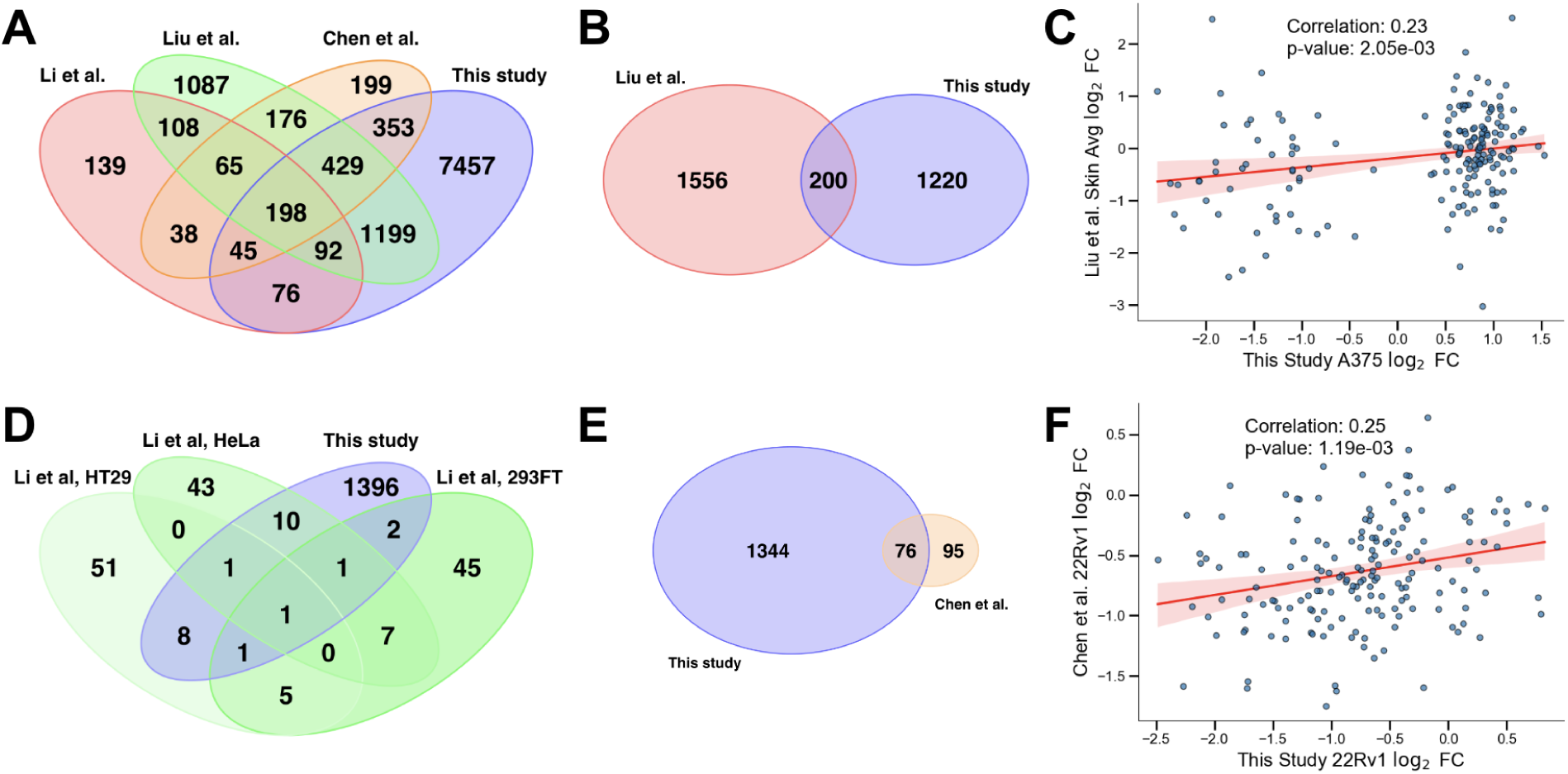
overlap of circRNAs screened in our study compared with previous studies. A) Venn diagram illustrating the overlap of circRNAs screened across four studies: Li et al. (red, 762 circRNAs), Liu et al. (green, 3,354 circRNAs), Chen et al. (orange, 1,507 circRNAs), and our study (blue, 9,853 circRNAs). The diagram shows that 429 circRNAs are shared among all four studies, while a significant number of circRNAs are unique to our study. B) Venn diagram showing the overlap of essential circRNAs identified in at least one cell line between Liu et al. (1,556 essential circRNAs) and our study (1,344 essential circRNAs), with 200 shared essential circRNAs. C) Scatter plot comparing circRNA log_2_ fold changes from the A375 melanoma cell line in our study (x-axis) and the average skin log_2_ fold change from Liu et al. (y-axis). (r = 0.23, p = 2.05 x 10^-3^) F) Scatter plot comparing circRNA log_2_ fold changes from the 22Rv1 prostate cancer cell line in our study (x-axis) and Chen et al.’s results (y-axis). (r = 0.25, p = 1.19 x 10^-3^)

**Supplementary Table 1:**

Comparisons final adjusted circRNA shRNA score and corresponding average log2 fold change from the screen

**Supplementary Table 2:**

Comparisons final adjusted circRNA shRNA score and corresponding T8vsT0 robust rank agreement score from the screen

**Supplementary Table 3:**

Comparison of log_2_ fold change across cell lines for linear transcripts, positive controls, and negative controls across T8 and T16 samples relative to T0.

**Supplementary Table 4:**

Comparison of dropout score across the screened cell lines for the union of essential linear transcripts and the DepMap shRNA screening project.

**Supplementary Table 5:**

Comparison of log_2_ fold change across cell lines for circular transcripts, positive controls, and negative controls across T8 and T16 samples relative to T0.

**Supplementary Table 6:**

Primer, oligo sequences and siRNAs used in this study.

## References

1. Slack, F. J. & Chinnaiyan, A. M. The Role of Non-coding RNAs in Oncology. Cell vol. 179 1033–1055 Preprint at 10.1016/j.cell.2019.10.017 (2019).

2. Esposito, R. et al. Hacking the Cancer Genome: Profiling Therapeutically Actionable Long Non-coding RNAs Using CRISPR-Cas9 Screening. Cancer Cell 35, 545–557 (2019).

3. Anfossi, S., Babayan, A., Pantel, K. & Calin, G. A. Clinical utility of circulating non-coding RNAs - an update. Nat. Rev. Clin. Oncol. 15, 541–563 (2018).

4. St Laurent, G., Wahlestedt, C. & Kapranov, P. The Landscape of long noncoding RNA classification. Trends Genet. 31, 239–251 (2015).

5. Chen, L.-L. The expanding regulatory mechanisms and cellular functions of circular RNAs. Nat. Rev. Mol. Cell Biol. 21, 475–490 (2020).

6. Liu, J., Zhang, X., Yan, M. & Li, H. Emerging Role of Circular RNAs in Cancer. Frontiers in Oncology vol. 10 Preprint at 10.3389/fonc.2020.00663 (2020).

7. Rajappa, A., Banerjee, S., Sharma, V. & Khandelia, P. Circular RNAs: Emerging Role in Cancer Diagnostics and Therapeutics. Front Mol Biosci 7, 577938 (2020).

8. Vo, J. N. et al. The Landscape of Circular RNA in Cancer. Cell 176, 869–881.e13 (2019).

9. Chen, S. et al. Widespread and Functional RNA Circularization in Localized Prostate Cancer. Cell 176, 831–843.e22 (2019).

10. Smid, M. et al. The circular RNome of primary breast cancer. Genome Res. 29, 356–366 (2019).

11. Zhang, Y. et al. Optimized RNA-targeting CRISPR/Cas13d technology outperforms shRNA in identifying functional circRNAs. Genome Biol. 22, 41 (2021).

12. Li, S. et al. Screening for functional circular RNAs using the CRISPR-Cas13 system. Nat. Methods 18, 51–59 (2021).

13. He, A. T., Liu, J., Li, F. & Yang, B. B. Targeting circular RNAs as a therapeutic approach: current strategies and challenges. Signal Transduct Target Ther 6, 185 (2021).

14. Chen, S., Zhang, J. & Zhao, F. Screening Linear and Circular RNA Transcripts from Stress Granules. Genomics Proteomics Bioinformatics (2022) doi:10.1016/j.gpb.2022.01.003.

15. Lee-Yow, Y. C., Valbuena, R. C., Richter, C. S., Chang, H. Y. & Engreitz, J. M. Junction-targeting designs limit the application of CRISPR-Cas13d in circular RNA perturbation studies. Nucleic Acids Res 54, (2026).

16. Hart, T. et al. High-Resolution CRISPR Screens Reveal Fitness Genes and Genotype-Specific Cancer Liabilities. Cell 163, 1515–1526 (2015).

17. Tsherniak, A. et al. Defining a Cancer Dependency Map. Cell 170, 564 (2017).

18. Nagpal, N. et al. Essential role of MED1 in the transcriptional regulation of ER-dependent oncogenic miRNAs in breast cancer. Sci. Rep. 8, (2018).

19. Seachrist, D. D., Anstine, L. J. & Keri, R. A. FOXA1: A Pioneer of Nuclear Receptor Action in Breast Cancer. Cancers 13, (2021).

20. Matevossian, A. & Resh, M. D. Hedgehog Acyltransferase as a target in estrogen receptor positive, HER2 amplified, and tamoxifen resistant breast cancer cells. Mol. Cancer 14, 1–15 (2015).

21. Brett, J. O., Spring, L. M., Bardia, A. & Wander, S. A. ESR1 mutation as an emerging clinical biomarker in metastatic hormone receptor-positive breast cancer. Breast Cancer Res. 23, 1–15 (2021).

22. del Mar Maldonado, M. & Dharmawardhane, S. Targeting Rac and Cdc42 GTPases in Cancer. Cancer Res. 78, 3101 (2018).

23. Xiao, X.-H. et al. Regulating Cdc42 and Its Signaling Pathways in Cancer: Small Molecules and MicroRNA as New Treatment Candidates. Molecules : A Journal of Synthetic Chemistry and Natural Product Chemistry 23, (2018).

24. Lee, Y. G. et al. LONP1 and ClpP cooperatively regulate mitochondrial proteostasis for cancer cell survival. Oncogenesis 10, 1–14 (2021).

25. Bedi, U. et al. SUPT6H controls estrogen receptor activity and cellular differentiation by multiple epigenomic mechanisms. Oncogene 34, (2015).

26. Su, W. et al. Overexpressed WDR3 induces the activation of Hippo pathway by interacting with GATA4 in pancreatic cancer. J. Exp. Clin. Cancer Res. 40, 1–15 (2021).

27. Wang, Y. et al. RPS24 knockdown inhibits colorectal cancer cell migration and proliferation in vitro. Gene 571, (2015).

28. Chen, H.-H. et al. Circular RNA detection identifies circPSEN1 alterations in brain specific to autosomal dominant Alzheimer’s disease. Acta Neuropathol Commun 10, 29 (2022).

29. Yao, S. et al. CircRNA ARFGEF1 functions as a ceRNA to promote oncogenic KSHV-encoded viral interferon regulatory factor induction of cell invasion and angiogenesis by upregulating glutaredoxin 3. PLoS Pathog. 17, e1009294 (2021).

30. Yu, D. & Zhang, C. Circular RNA PTK2 Accelerates Cell Proliferation and Inhibits Cell Apoptosis in Gastric Carcinoma via miR-139-3p. Dig. Dis. Sci. 66, 1499–1509 (2021).

31. Zhou, F. et al. Circular RNA Protein Tyrosine Kinase 2 Promotes Cell Proliferation, Migration and Suppresses Apoptosis via Activating MicroRNA-638 Mediated MEK/ERK, WNT/β-Catenin Signaling Pathways in Multiple Myeloma. Front. Oncol. 11, 648189 (2021).

32. Ma, L. et al. Silencing of circRACGAP1 sensitizes gastric cancer cells to apatinib via modulating autophagy by targeting miR-3657 and ATG7. Cell Death Dis. 11, 169 (2020).

33. Zhou, Z. et al. Circular RNAs act as regulators of autophagy in cancer. Mol Ther Oncolytics 21, 242–254 (2021).

34. Yang, Q. et al. Circular RNA expression profiles during the differentiation of mouse neural stem cells. BMC Syst. Biol. 12, 128 (2018).

35. Jiang, Z. et al. Circular RNA protein tyrosine kinase 2 (circPTK2) promotes colorectal cancer proliferation, migration, invasion and chemoresistance. Bioengineered 13, 810–823 (2022).

36. Liu, L. et al. Systematic loss-of-function screens identify pathway-specific functional circular RNAs. Nat. Cell Biol. 26, 1359–1372 (2024).

37. Li, J. et al. A genomic and epigenomic atlas of prostate cancer in Asian populations. Nature 580, 93–99 (2020).

38. Fraser, M. et al. Genomic hallmarks of localized, non-indolent prostate cancer. Nature 541, 359–364 (2017).

39. Liu, S. J. et al. CRISPRi-based genome-scale identification of functional long noncoding RNA loci in human cells. Science 355, (2017).

40. Liu, X. et al. Circular RNA: An emerging frontier in RNA therapeutic targets, RNA therapeutics, and mRNA vaccines. J. Control. Release 348, 84–94 (2022).

41. Misir, S., Wu, N. & Yang, B. B. Specific expression and functions of circular RNAs. Cell Death Differ. 29, 481–491 (2022).

42. Cheng, Q. et al. Elevated MPP6 expression correlates with an unfavorable prognosis, angiogenesis and immune evasion in hepatocellular carcinoma. Front. Immunol. 14, 1173848 (2023).

43. Xu, F. et al. Downregulating SynCAM and MPP6 expression is associated with ovarian cancer progression. Oncol. Lett. 18, 2477–2483 (2019).

44. Ai, Y., Liang, D. & Wilusz, J. E. CRISPR/Cas13 effectors have differing extents of off-target effects that limit their utility in eukaryotic cells. Nucleic Acids Res. 50, e65 (2022).

45. Bayoumi, M. & Munir, M. Potential Use of CRISPR/Cas13 Machinery in Understanding Virus-Host Interaction. Front. Microbiol. 12, 743580 (2021).

46. Glažar, P., Papavasileiou, P. & Rajewsky, N. circBase: a database for circular RNAs. RNA 20, 1666–1670 (2014).

47. Zhang, X.-O. et al. Diverse alternative back-splicing and alternative splicing landscape of circular RNAs. Genome Res. 26, 1277–1287 (2016).

48. Chen, X. et al. circRNADb: A comprehensive database for human circular RNAs with protein-coding annotations. Sci. Rep. 6, 34985 (2016).

49. Xia, S. et al. CSCD: a database for cancer-specific circular RNAs. Nucleic Acids Res. 46, D925–D929 (2018).

50. Li, W. et al. MAGeCK enables robust identification of essential genes from genome-scale CRISPR/Cas9 knockout screens. Genome Biol. 15, 554 (2014).

51. Dobin, A. et al. STAR: ultrafast universal RNA-seq aligner. Bioinformatics 29, 15–21 (2013).

52. Wang, L., Wang, S. & Li, W. RSeQC: quality control of RNA-seq experiments. Bioinformatics 28, 2184–2185 (2012).

53. Wang, L. et al. Measure transcript integrity using RNA-seq data. BMC Bioinformatics 17, 58 (2016).

54. Anders, S., Pyl, P. T. & Huber, W. HTSeq--a Python framework to work with high-throughput sequencing data. Bioinformatics 31, 166–169 (2015).

55. Love, M. I., Huber, W. & Anders, S. Moderated estimation of fold change and dispersion for RNA-seq data with DESeq2. Genome Biol. 15, 550 (2014).

56. Wickham, H. ggplot2: Elegant Graphics for Data Analysis. (Springer, 2016).

57. Wang, B. et al. Integrative analysis of pooled CRISPR genetic screens using MAGeCKFlute. Nat. Protoc. 14, 756–780 (2019).

58. Chen, H. & Boutros, P. C. VennDiagram: a package for the generation of highly-customizable Venn and Euler diagrams in R. BMC Bioinformatics 12, 35 (2011).

59. Lex, A., Gehlenborg, N., Strobelt, H., Vuillemot, R. & Pfister, H. UpSet: Visualization of Intersecting Sets. IEEE Trans. Vis. Comput. Graph. 20, 1983–1992 (2014).

60. Gu, Z., Eils, R. & Schlesner, M. Complex heatmaps reveal patterns and correlations in multidimensional genomic data. Bioinformatics 32, 2847–2849 (2016).

61. P’ng, C. et al. BPG: Seamless, automated and interactive visualization of scientific data. BMC Bioinformatics 20, 42 (2019).

62. Cui, X. et al. Guitar: An R/Bioconductor Package for Gene Annotation Guided Transcriptomic Analysis of RNA-Related Genomic Features. Biomed Res. Int. 2016, 8367534 (2016).

63. Wu, T. et al. clusterProfiler 4.0: A universal enrichment tool for interpreting omics data. Innovation (Camb) 2, 100141 (2021).

64. AnnotationDbi. Bioconductor https://bioconductor.org/packages/AnnotationDbi.

65. Yu, G., Wang, L.-G., Yan, G.-R. & He, Q.-Y. DOSE: an R/Bioconductor package for disease ontology semantic and enrichment analysis. Bioinformatics 31, 608–609 (2015).

66. Carlson, M. *org.Hs.eg.db*. (Bioconductor, 2017). doi:10.18129/B9.BIOC.ORG.HS.EG.DB.

67. Blighe, K. EnhancedVolcano. (Bioconductor, 2018). doi:10.18129/B9.BIOC.ENHANCEDVOLCANO.

68. Waskom, M. seaborn: statistical data visualization. J. Open Source Softw. 6, 3021 (2021).

69. Virtanen, P. et al. SciPy 1.0: fundamental algorithms for scientific computing in Python. Nat. Methods 17, 261–272 (2020).

70. Hunter, J. D. Matplotlib: A 2D Graphics Environment. Comput. Sci. Eng. 9, 90–95 (May-June 2007).

71. https://cran.r-project.org/web/packages/shiny/index.html.

