## Supplementary Table 1 to 5 for "Functional landscape of circular RNAs in human cancer cells"

**Supplementary Table 1: Comparisons final adjusted circRNA shRNA score and corresponding average log_2_ fold change from the screen**

| **Cell line** | **Correlation** | **p-value** |
| --- | --- | --- |
| 22Rv1 | -0.013 | 0.2076 |
| A375 | -0.084 | < 0.001 |
| MCF7 | -0.056 | 4.58x10 ^-6^ |
| HCT116 | -0.061 | 1.32x10 ^-9^ |
| FTC133 | -0.031 | 0.0022 |
| HeLa | -0.034 | 0.0007 |

**Supplementary Table 2: Comparisons final adjusted circRNA shRNA score and corresponding T8vsT0 robust rank agreement score from the screen**

| **Cell line** | **Correlation** | **p-value** |
| --- | --- | --- |
| 22Rv1 | -0.011 | 0.282 |
| A375 | -0.078 | 1.55x10 ^-14^ |
| MCF7 | -0.043 | 2.76x10 ^-5^ |
| HCT116 | -0.063 | 7.20x10 ^-10^ |
| FTC133 | -0.013 | 0.1897 |
| HeLa | -0.029 | 0.0042 |

**Supplementary Table 3: Comparisons for T8 and T16 samples relative to T0 separately for linear screen**

| **Cell line** | **Correlation** | **p-value** |
| --- | --- | --- |
| A549 | 0.83 | < 0.001 |
| A375 | 0.85 | < 0.001 |
| MCF7 | 0.78 | < 0.001 |
| HCT116 | 0.91 | < 0.001 |
| FTC133 | 0.61 | < 0.001 |
| HeLa | 0.74 | < 0.001 |

**Supplementary Table 4: Comparisons for essential and non-essential genes identified by the DepMap project for each cell line**

| **Cell line** | **Correlation** | **p-value** |
| --- | --- | --- |
| A549 | 0.54 | 3.10 x 10^-9^ |
| A375 | 0.52 | 5.37 x 10^-8^ |
| MCF7 | 0.60 | 7.12 x 10^-10^ |
| HCT116 | 0.64 | 4.75 x 10^-15^ |
| FTC133 | 0.56 | 6.26 x 10^-8^ |
| HeLa | 0.38 | 7.27 x 10^-5^ |

**Supplementary Table 5: Comparisons for T8 and T16 samples relative to T0 separately for circular screen**

| **Cell line** | **Correlation** | **p-value** |
| --- | --- | --- |
| A549 | 0.87 | < 0.001 |
| A375 | 0.88 | < 0.001 |
| MCF7 | 0.80 | < 0.001 |
| HCT116 | 0.93 | < 0.001 |
| FTC133 | 0.63 | < 0.001 |
| HeLa | 0.80 | < 0.001 |
